# Sporulation modulates viscoelastic development through extracellular matrix restructuring in *Bacillus subtilis* biofilms

**DOI:** 10.64898/2026.09.23.753867

**Authors:** Nicolas A. Ducharme, Joseph W. Larkin

## Abstract

Biofilm formation–the establishment of cellular communities within a self-secreted polymer matrix–is a behavior exhibited by almost all species of bacteria. Bacterial biofilms are well described as viscoelastic materials, with properties that grant physical advantages, such as elastic resilience and viscous adaptability. Additionally, many biofilm-forming species produce a subpopulation of highly resilient spores. In the model biofilm- and spore-forming species *Bacillus subtilis*, sporulation and matrix production are regulated by a common gene pathway, provoking the question of how sporulation influences biofilm viscoelastic properties. Investigating this point will facilitate the management of both salutary and deleterious biofilms, especially through manipulating their establishment and dispersion. Here, we investigate the interplay of sporulation and viscoelastic properties using rheological measurements of *B. subtilis* biofilms with varied matrix and spore production. We find that, to a significantly greater extent than matrix production, biofilm physical development is strongly altered by activation of sporulation through the *spo0A* regulation pathway. Spore-deficient strains initially establish physically robust biofilms with linear viscoelastic properties and recovery behavior distinct from spore-producing biofilms. However, spore-deficient biofilms fail to maintain their physical properties over time. In contrast, spore-producing wild-type and matrix-mutant biofilms exhibit more stable physical properties. Our results demonstrate that cell-level gene regulation is associated with macroscopic mechanical transitions in cellular communities, with sporulation granting not only a cellular-level survival advantage, but also greater community-level physical stability.

**SIGNIFICANCE:** Biofilms are living materials with physical properties that influence their resilience. It is known that extracellular matrix composition largely determines biofilm physical properties. However, the role of cellular development pathways on biofilm extracellular matrix remains underexplored. Here, we examine the role of the cellular development pathway of sporulation–the production of resilient spores–in the development of biofilm physical properties. We find that spore-producing biofilms are initially softer and store less stress than spore-deficient biofilms. However, spore-producing biofilms maintain their physical properties over time whereas spore-deficient biofilms become softer and recover more poorly. Our results suggest sporulation benefits biofilms by maintaining their physical properties at the cost of decreased stiffness, which has important implications for their survival and persistence.

## INTRODUCTION

Bacteria commonly form living viscoelastic materials called biofilms (1–3) whose physical properties play key roles throughout their development and life cycle, mediating their survival and propagation (4) from the initial adhesion of cell aggregates to surfaces (5) to the dispersion of cells from mature biofilms (6). Bacteria are capable of sensing environmental forces and adjusting their physical properties in response (7, 8), thereby providing bacterial cells protection against environmental perturbations (9), resistance to fluid flow and deformation (4, 9, 10), and recovery from damage (9). These physical advantages are important in the dispersal and persistence of biomass (11), including in clinical and industrial settings (2, 12). Due to the ubiquity of biofilms and their impact on human well-being, investigating the links between microscopic structure and macroscopic mechanical response in biofilms is a pursuit of great interest and importance (13). Understanding the physical properties and processes that govern biofilm survival is crucial to modeling biofilm resilience and developing microbial management techniques.

The extracellular matrix (ECM) that bacteria produce is key in determining biofilm physical attributes (14). Numerous factors influence ECM properties, including polymeric composition (15), network connectivity and architecture (16), and effective cross-link type and density (17). These factors also impact biofilm structure (3) which directly correlates with biofilm viscoelastic properties (18). An advantage bacteria gain from ECM production is influencing the localization of cell death which contributes to biofilm wrinkling and 3D structure, thereby providing localized stress relief (19). Bacteria further leverage ECM properties and structure to their benefit, collectively modifying matrix gene expression in response to external stressors, such as antibiotics, thereby enhancing biofilm formation (20).

In addition to modifying matrix gene expression, many bacterial species respond to the stress of nutrient depletion and high cell density (21–23) by producing metabolically dormant spores that are highly resilient to many environmental conditions (24). Although advantageous to the bacteria, spore dispersal is problematic for humans, being a key mechanism for the spread of food-borne pathogens (25). The broadly conserved transcription factor Spo0A (26) regulates both sporulation and matrix production in many spore-forming species (27–29), genetically linking biofilm viscoelastic properties and sporulation. This genetic link between sporulation and matrix production has important effects on the biofilm life cycle, with biofilm physical properties impacting the dispersal of spores from mature biofilms (30). There are numerous additional ways in which the developmental switch from vegetative cell to dormant spore could impact emergent biofilm properties. First, spores are metabolically inactive and therefore neither consume nutrients nor produce ECM components (31), thereby potentially affecting the nutrient concentration and ECM density. Second, when the mother cell lyses during sporulation, releasing the spore along with other cellular contents into the ECM (32, 33) could disrupt cell-ECM connections and the biofilm polymer network. Third, the ECM may also potentially interact with spores directly, integrating them into the polymer network. Despite the close feedback between sporulation, biofilm material properties, and spore dispersal, the relationship between sporulation and biofilm viscoelasticity has been minimally studied. Investigating their interplay will have important implications for managing biofilms and spore dispersal (25).

Here we determine the relationship between sporulation and biofilm viscoelasticity by performing rheological measurements on the soil bacterium *Bacillus subtilis. B. subtilis* is an ideal model organism to investigate the effect of sporulation on biofilm viscoelastic properties: *B. subtilis* has been thoroughly studied as both a biofilm- and spore-former (34–36), and the genetic link between matrix production and sporulation is well understood. A protein network called the phosphorelay senses environmental and cell density conditions and ultimately phosphorylates Spo0A. The concentration of phosphorylated Spo0A is crucial in determining cell fate, with intermediate levels activating matrix production and high levels activating sporulation (27). Using genetic knockouts, we find that sporulation has a greater impact on the development of biofilm viscoelastic properties than the production of two key ECM polymers, exopolysaccharides (EPS) (37) and poly-*γ*-glutamic acid (PGA) (38), despite the central roles of these polymers in biofilm integrity and structure (39, 40).

We propose that sporulation increases the extent of polymer network restructuring, modifying the development of the biofilm physical stress response over time–a hypothesis with which our data is consistent. Our findings provide a basis for establishing causal links between developmental processes, matrix composition, and bulk viscoelastic properties, the expanded understanding of which will will facilitate determining underlying biofilm polymer network properties using bulk rheological measurements.

## MATERIALS AND METHODS

### Bacterial strains

In this work, we used the base strain of *Bacillus subtilis* NCIB3610 (41) with deletion of the plasmid-borne gene *rapP* and constitutive expression of mScarlet via the construct *P*_*veg*_-*mScarlet*. The deletion of *rapP* restores biofilm sporulation levels to those observed in environmental isolates (42). We refer to this strain as wild-type (WT) throughout the text. From this strain, we constructed three mutants: a poly-*γ*-glutamic acid (PGA) knockout via deletion of the *pgsBCAE* operon, an exopolysaccharide (EPS) knockout via deletion of the *epsAO* operon, and a spore knockout via deletion of the *spo0A* promoter *P*_*s*_. The stationary phase promoter *P*_*s*_ becomes active during starvation to initiate sporulation. Deletion of *P*_*s*_ results in a strain that can form robust biofilms that contain no mature spores (43). Refer to Table 1 for a summary of the strains used herein.

**Table 1.** Bacterial strains. Designations, genotypes, and sources for the Bacillus subtilis strains used in this work.

| Strain Designation | Relevant Genotype | Source, (reference) |
| --- | --- | --- |
| JMJ851 | 3610 | BGSC code 3A38, (41) |
| JMJ1155 | 3610 $\Delta rapP::tet$ | Jones, (42) |
| JMJ1233 (WT) | 3610 $\Delta rapP::tet lacA::Pveg-mScarlet-kan$ | Jones, (42) |
| JMJ1234 (Spore-) | 3610 $\Delta rapP::tet lacA::Pveg-mScarlet-kan spo0A\Delta Ps$ | Jones, (42) |
| JMJ1250 (PGA-) | 3610 $\Delta rapP::tet lacA::Pveg-mScarlet-kan \Delta pgsB::cat$ | Jones, (42) |
| JMJ1480 (EPS-) | 3610 $\Delta rapP::tet \Delta epsA-O::tet$ | Jones, (42) |

### Plates and media

For all experiments, we grew biofilms in 100 mm Petri dishes on 2% agar with Minimal Salts glycerol glutamate (MSgg) media adapted from Branda et al. (44) (0.5% w/v monosodium glutamate, 0.5% v/v glycerol,100 mM MOPS (pH 7.0), 5 mM potassium phosphate (pH 7.0), 700 μM CaCl_2_, 2 mM MgCl_2_, 50 μM MnCl_2_, 100 μM FeCl_3_ 6H_2_O, 1 μM ZnCl_2_,2 μM thiamine HCl). We prepared cultures by isolating single colonies from lysogeny broth (LB) plates and growing them in liquid LB at 37°C to an OD_600_ of 0.7. We distributed 50 μL of these cultures on MSgg plates via sterile glass beads, then incubated them at 30°C for 24 or 72 hours, resulting in lawn biofilms with sufficient biomass for rheometer loading.

### Rheological assays

For rheological data collection, we used a TA Instruments DHR-2 Rheometer. For all rheological measurements, we used a 20 mm, 2° cone and plate geometry. We first brought the plate to 30°C using a Standard Peltier Plate Temperature Controller, following which we zeroed the plate gap. We then extracted the biomass from the agar plates via gentle scraping parallel to the surface with a glass slide and transferred it to the rheometer plate (Fig. 2A). Next, we adjusted the plate gap to 83 μm and trimmed excess biomass using a glass slide. We then enclosed the geometry in a hood wrapped with Parafilm (to mitigate runtime desiccation) and set the gap to 62 μm. We performed all rheological assays, with the exception of the frequency sweep, in oscillatory mode with an angular frequency of 10.0 rad/s. The fundamental quantities determined by oscillatory rheology are the shear storage and loss moduli, with the storage modulus corresponding to the material’s stiffness and the loss modulus its viscosity.

**Figure 1.**
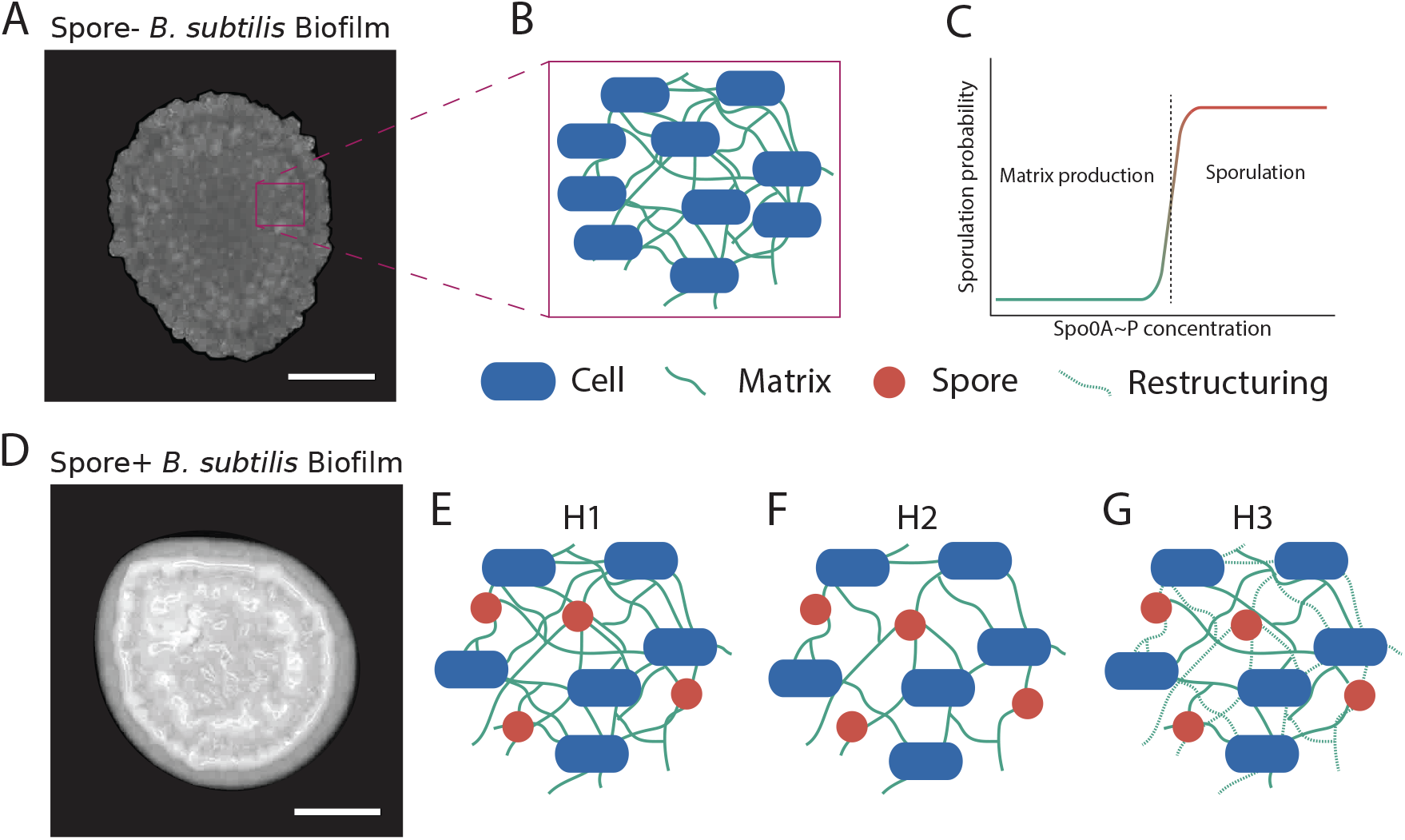
Sporulation-associated changes in biofilm polymer network may explain qualitative property changes. (A) Stereo microscope image of Δ*P*_*s*_ (spore-) *Bacillus subtilis* biofilm at 24 hours (Scale bar: 1 mm). (B) Illustration of spore-knockout biofilm polymer network, with vegetative cells and matrix components. (C) Conceptual plot of sporulation probability vs Spo0A∼P concentration. When the concentration of Spo0A∼P are below the threshold for sporulation, matrix genes can be expressed. (D) Stereo microscope image of WT (spore+) *Bacillus subtilis* biofilm at 24 hours (Scale bar: 1 mm). The spore+ biofilm (D) differs from the spore-biofilm (A) in its qualitative features, including glossiness, edge roughness, and surface topology. The exposure settings are identical for (A) and (D), so the difference in pixel intensity indicates a difference in reflectivity. (E) Illustration of spore-producing biofilm polymer network under the “Rigid Inclusions” hypothesis, **H1**. Compared to (B), all spores are replaced by vegetative cells. (F) The “Network Density” hypothesis, **H2**, in which the smaller proportion of vegetative cells leads to a lower polymer network density. (G) The “Network Restructuring” hypothesis, **H3**, in which sporulation alters polymer network restructuring processes, with “Restructuring” referring to these modified processes collectively.

**Figure 2.**
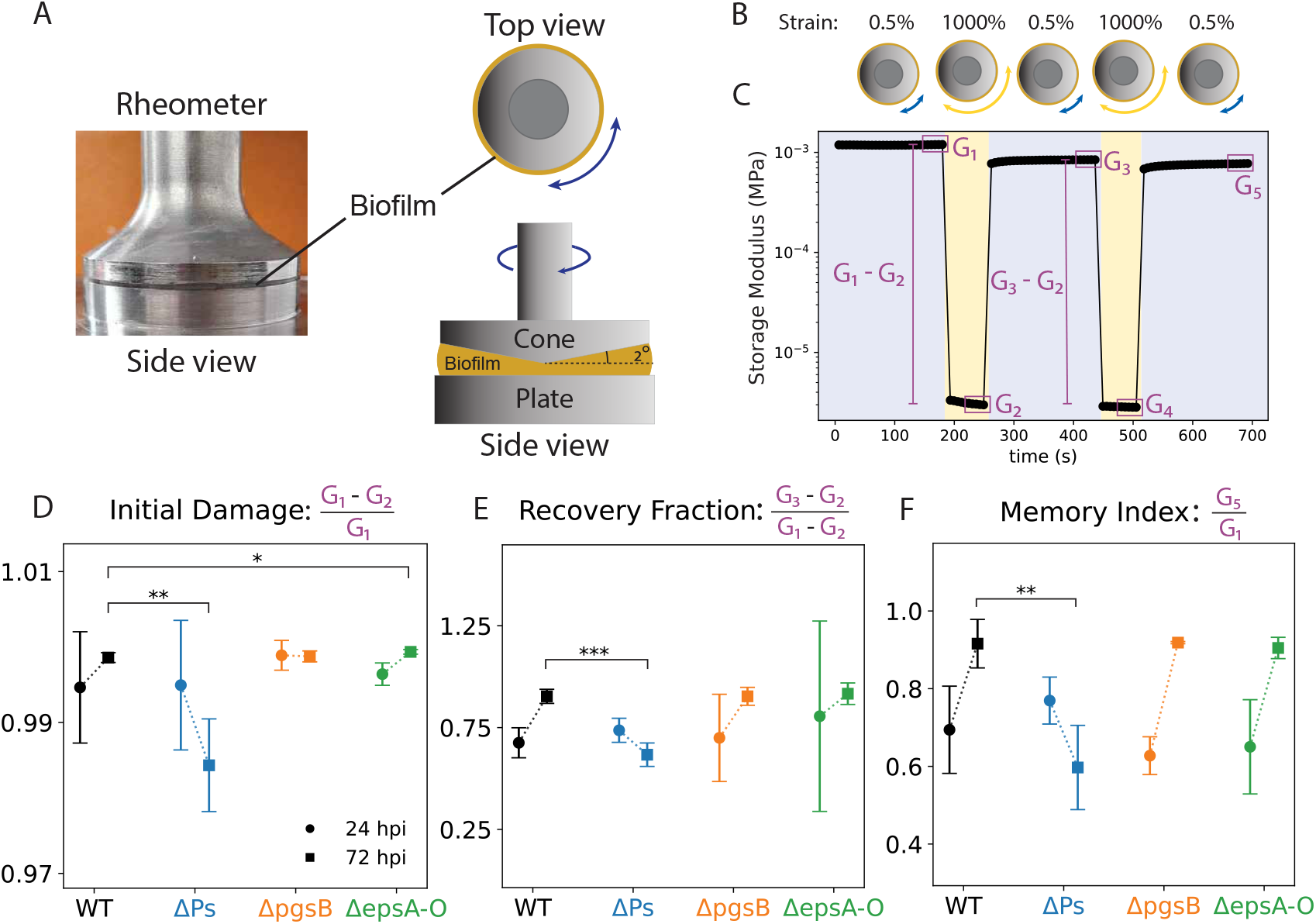
Sporulation associated with consistent trends in biofilm recovery following high strain. (A) Side-view photo (left) and illustrations (right) of a rheometer with biofilm between the cone and plate. (B) Top-down illustration of rheometer alternating between periods of low-amplitude (blue) and high-amplitude (yellow) strain. (C) A representative dataset from the rheological recovery protocol, with periods of low-amplitude strain shaded in blue and high-amplitude strain in yellow. The means of the boxed data constitute G_1_ through G_5_. (D) Mean initial damage. The 72-hour spore-knockout biofilms have a mean 98% that of WT. At 72 hours, the EPS-knockout biofilms show a statistically, but not physically, significant difference from WT. (E)Mean recovery fraction. At 72 hours, the spore-knockout biofilms have a mean 68.9% that of WT. The spore-producing biofilms increase in average recovery fraction from 24 to 72 hours, with WT increasing by 28.6% of its 24-hour value. (F) Mean memory index. The 72-hour spore-knockout biofilms have a mean 66% that of WT. Spore-knockout biofilms have a memory index 80% of their 24-hour mean, while spore-producing biofilms have means that increase between 29% and 39%. (D) - (F) Data collected from n = 3 independent samples. Error bars represent 95% confidence intervals determined by the Margin of Error (ME = SEM × t^∗^). *p<0.05, **p<0.01, ***p<0.001. P-values calculated by Welch’s two-tailed t-test on independent samples.

The recovery assay quantifies the degree to which biofilms recover their stiffness following high strain. In this assay, we alternated the strain amplitude between 0.5% and 1000%. With a 2° rheometer cone, a strain of *p*% corresponds to an angular displacement of 2 [ineqq] so 0.5% and 1000% strains are equivalent to angular displacements of 0.01° and 20°, respectively. In this assay, we set the strain amplitude to 0.5% for 180 s, then 1000% for 60 s (Fig. 2BC). We repeated this cycle once, following which we set the amplitude to 0.5% for 180 s. We calculated the initial damage, recovery fraction, and memory index by comparing the storage moduli during the phases of this cycle (Fig. 2D-F).

Our second rheological assay, the frequency sweep, quantifies the dependence of the storage (*G*^′^) and loss (*G*^″^) moduli on processes active at different time scales. We performed this assay by sweeping from 1.0 to 100.0 rad/s, holding the oscillation amplitude at 0.5%.

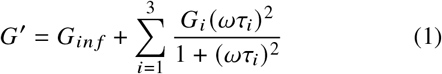

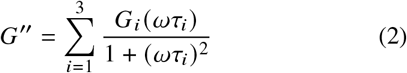

We fit the data to the most general linear model of viscoelasticity, the generalized Maxwell model, with three modes (Eqs. 1-2) (Fig. 3A). Here, *G*_*inf*_ is the non-relaxing modulus, *G*_*i*_ are the mode moduli, *ω* is the angular frequency, and τ_*i*_ are the relaxation times. We fixed the relaxation times τ_*i*_ to 0.01, 0.1, and 1.0 seconds, based approximately on the range of frequencies used in this assay (1.0 to 100.0 rad/s), and fit the corresponding mode moduli *G*_*i*_ and *G*_*inf*_ to the data (Fig. 3B). We also fit the data to one- and two-mode Maxwell models with poor results (see Fig. S4). The three-mode model was the lowest-degree model capable of recapitulating the data (see fit residuals in Fig. S4).

**Figure 3.**
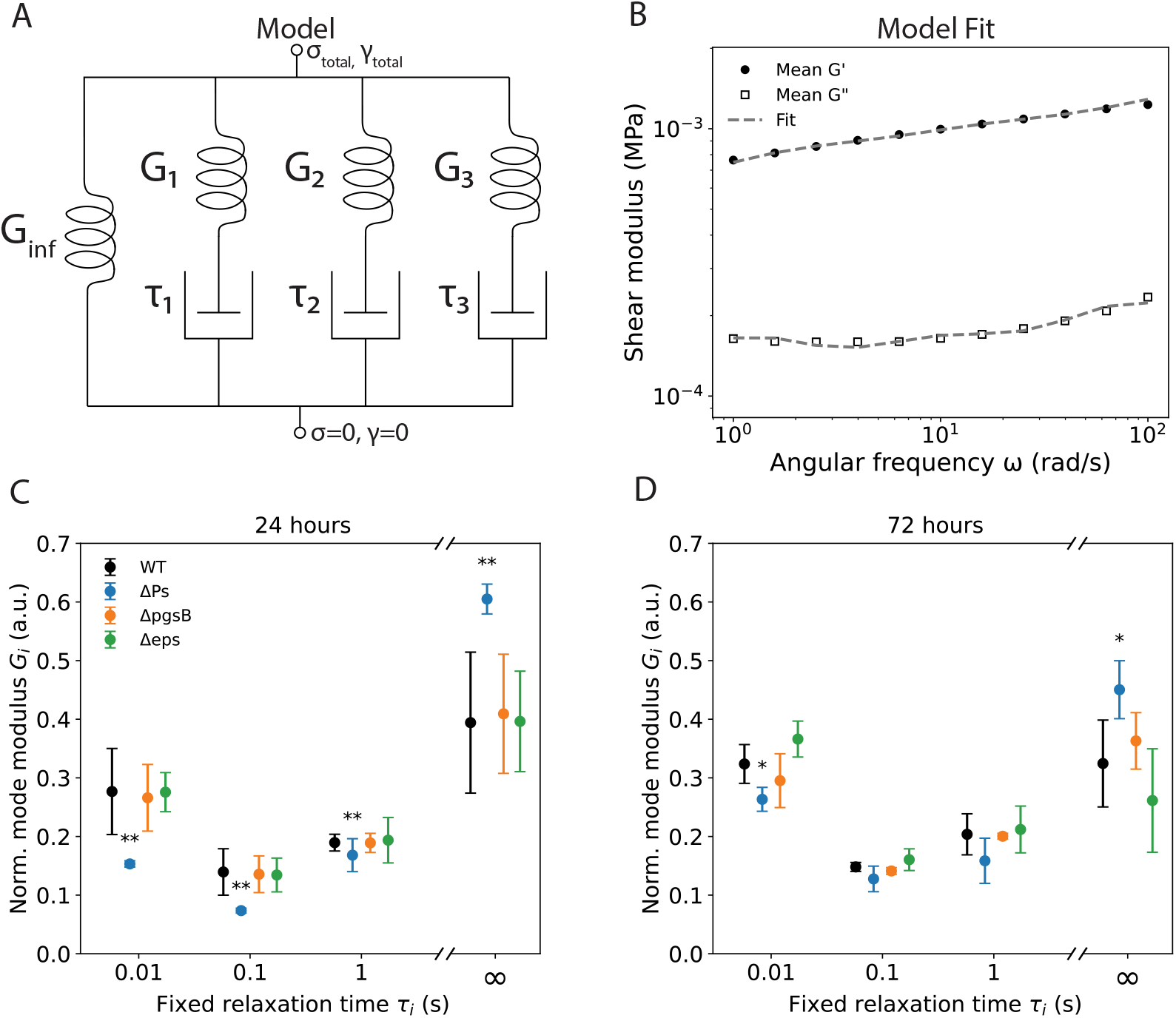
Sporulation narrows and maintains biofilm relaxation spectrum. (A) Schematic of three-mode generalized Maxwell model. Spring moduli are denoted by G and dashpot relaxation times by τ. The stress and strain are denoted by *σ* and *γ*, respectively. (B) Representative WT frequency sweep data, averaged from n = 3 independent samples, with Maxwell model fit. The fit was performed using ordinary least squares. (C) Plot of normalized mode modulus vs relaxation time for 24-hour biofilms. All three normalized mean mode moduli are lower for spore-knockout biofilms than spore-producing biofilms. The 0.01 s norm. modulus is 55% of WT. The 0.1 s norm. modulus is 54% of WT. The 1 s norm. modulus is 88% of WT. The spore-knockout biofilms has an average *G*_*inf*_ 1.5× WT. (D) Plot of normalized mode modulus vs relaxation time for 72-hour biofilms. The spore-knockout biofilms have a lower normalized mean mode modulus corresponding to τ = 0.01 s, being 80% of WT. The spore-knockout biofilms had an average *G*_*inf*_ 1.5× WT. (C)- (D) Fits were performed onn=3 independent samples. The fitting parameters for each replicate were normalized by dividing by their sum (including G_*inf*_). Error bars represent 95% confidence intervals determined by the Margin of Error (ME = SEM × t^∗^). *p<0.05, **p<0.01. P-values compare data to WT at the same time point and relaxation time and were calculated by Welch’s two-tailed t-test on independent samples.

Our third and final rheological assay, the strain sweep, quantifies the dependence of the storage and loss moduli on the magnitude of the strain. In this assay, we swept the oscillation strain amplitude from 0.01% to 200%, recording the biofilm’s storage and loss moduli. From these data, we calculated the plateau moduli, which are the moduli under low strain conditions. We calculated the plateau moduli as the median of the three lowest-strain data points. The loss factor is the ratio of the loss modulus to the storage modulus. When the loss factor is between 0.0 and 1.0, the material is more solid-like, and when it is greater than 1.0 it is more liquid-like.

### Stereo microscope images

We used an Olympus SZX7 Zoom Stereo Microscope to image micro-colonies (Fig. 1AD) and record videos (see Videos S1-S4 at https://github.com/Larkin-Lab/ducharme_larkin_sporulation.git) while perturbing them with the bottom of a micro-centrifuge tube.

### Vegetative cell and spore counts

To quantify the number of vegetative cells and spores present in biofilms, we performed colony-forming unit (CFU) counts (see Fig. S1). We performed total CFU counts (vegetative cells and spores) by transferring the biomass from a 100 mm Petri dish to a 15 mL conical tube. After thoroughly macerating the biomass using a sterile wooden stick, we added 10 mL of 1×PBS to the tube. We then scraped the inside of the tube with the wooden stick to dislodge and break apart biomass. Next, we capped and manually agitated the tube until no precipitate remained. We prepared a sterile 96-well plate with 225 μL of 1×PBS to which we added 25 μL from the tube (for a 10×dilution). We repeated this dilution process to produce 10^6^×and 10^7^×dilutions. We evenly distributed 100 μL of these dilutions on LB agar plates using sterile glass beads. We then incubated the plates overnight at 37°C before finally counting the resultant colonies.

We performed spore counts by sealing the 96-well plate from the total CFU assay with a Breathe-Easy^®^ membrane (Diversified Biotech BEM-1) and baking it at 85°C for 30 minutes. We then removed the membrane and plated 100 μL of the 10^6^× and 10^7^ × dilutions on LB agar plates, using sterile glass beads for distribution. After incubating the plates overnight at 37°C, we counted the resultant colonies.

We observed some variation in total CFU counts and spore percentages (see Fig. S1) though they largely did not correlate with rheological properties.

### Mass measurements

We performed mass measurements to determine the fractions of the biofilm mass that were from extracellular matrix (ECM) and bound water (see Fig. S2). We note that it has been reported that the ECM accounts for over 90% of biofilm dry mass (51), so the dry mass and ECM mass are nearly synonymous.

We performed mass measurements by scraping biomass from a plate and transferring it to a pre-weighed piece of aluminum foil. We measured the wet mass of the sample using an Entris II Essential Line Precision Balance. We then baked the sample at 85°C for 48 hours. Next, we brought the sample to room temperature inside a desiccation chamber, following which we measured the dry mass of the sample using the precision balance.

We found that rheological properties did not correlate with mass measurements (see Fig. S2).

## RESULTS

### Sporulation qualitatively modifies biofilms

To observe the effect of spore regulation on biofilm development, we grew WT and Δ*P*_*s*_ (spore-deficient) biofilms on MSgg for 24 hours and imaged them with a stereo microscope. We observed qualitative differences in surface and edge morphologies as well as glossiness between the WT and spore-deficient biofilms (Fig. 1AD). Biofilm material properties have an important effect on biofilm morphology (40, 52), so these qualitative differences in biofilm morphology suggest changes in the underlying biofilm polymer networks associated with sporulation within biofilms.

As a qualitative test of the physical properties of the biofilms, we displaced a small quantity of biomass from lawn biofilms using the bottom of a 1.5 mL microcentrifuge tube and observed their viscoelastic response using a stereo microscope (see Videos S1-S4 at https://github.com/Larkin-Lab/ducharme_larkin_sporulation.git). We observed that at 24 hours, WT biofilms showed greater recoil compared to Δ*P*_*s*_ biofilms. Additionally, spore-producing biofilms were more easily disrupted than spore-knockout biofilms. At 72 hours, often considered a maturation point for *B. subtilis* biofilms (53), the spore-producing biofilms continued to exhibit enhanced recoil, however the spore-deficient biofilm was as easily disrupted as the spore-producing biofilm. These results suggest that spores and/or spore regulation contributes to the viscoelastic properties of biofilms. Furthermore, spore-deficient biofilms underwent a shift in their viscoelastic properties from 24 to 72 hours, unlike the spore-producing biofilms, which maintained their response to mechanical perturbation. Our qualitative experiments suggest differences in viscoelastic properties between biofilms based on the capacity for sporulation that could not be accounted for by cell counts or mass measurements (see Supplemental Information (SI), Section 2).

We propose three hypotheses (**H1**-**H3**) for the impact of sporulation on biofilm viscoelastic properties, including the aforementioned qualitative differences. We made predictions for each of these hypotheses using simple theoretical models (see SI, Section 1); however, more complex models using a combination of these and other mechanisms are likely applicable as well. Nevertheless, even without specific molecular-level information on the structure of biofilm polymers, our simple models provide a basis for understanding our macroscopic rheological data.

Motivating our first hypothesis, it has been demonstrated that inclusions, such as cells, suppress stiffening in biopolymer networks under shear deformation (54), suggesting that **H1**: spores affect polymer network properties by acting as rigid inclusions. We chose the rule of mixtures to address this hypothesis, although this should not be interpreted as the only applicable model for this hypothesis. According to the rule of mixtures, the greater rigidity of spores relative to vegetative cells and matrix would result in biofilm stiffness and viscosity scaling with spore fraction (Fig. 1BE) (see SI, Section 1A).

Our second hypothesis was **H2**: sporulation, which decreases the number of vegetative cells, in turn decreases the quantity of ECM produced. According to polymer gel theory, near the sol-gel percolation point–which *B. subtilis* biofilms are known to be (40)–a lower network density would result in a softer, less-dense biofilm (Fig. 1BF) (see SI, Section 1B).

Our final hypothesis was **H3**: sporulation modifies the restructuring processes active in the biofilm. In soft glassy rheology, all mechanisms that modify elements of the polymer network (i.e., all processes that restructure the network) are subsumed into a mean-field (i.e., coarse-grained) parameter *x* known as the noise temperature (55). In the case that sporulation restructures the polymer network, soft glassy rheology would predict a modification of the mean-field parameter *x*, which would correspond to changes in the development of biofilm rheological properties (Fig. 1BG) (see SI, Section 1C).

Each hypothesis predicted different rheological biofilm properties, specifically biofilm response to high strain, the temporal spectrum of its relaxation, and its viscoelasticity. These three hypotheses, along with rheological property predictions for each, are found in Table 2 (see also SI, Section 1). To determine the degree to which sporulation alters viscoelasticity and which of our hypotheses (**H1**-**H3**, Fig. 1EFG, Table 2) best accounts for the effect of sporulation on biofilm material properties, we required quantitative measurements of those properties, for which we turned to rheology.

**Table 2.** Viscoelastic property hypotheses & predictions. Each row indicates the viscoelastic property change predicted by hypotheses **H1** - **H3** when the spore fraction is increased, with “-” indicating no change.

| Viscoelastic Property | **H1**: Rigid Inclusions<sup>a</sup> | **H2**: Network Density | **H3**: Network Restructuring<sup>d</sup> |
| --- | --- | --- | --- |
| Damage | - | - | Increases |
| Recovery | - | - | Increases/Decreases |
| Relaxation Spectrum | - | Broadens<sup>b</sup> | Narrows |
| Stiffness | Increases | Decreases<sup>c</sup> | Decreases |
| Viscosity | - | Increases<sup>c</sup> | Decreases |

<sup>a</sup> Refs. (45, 46); <sup>b</sup> Refs. (47, 48); <sup>c</sup> Ref. (49); <sup>d</sup> Ref. (50)

### Sporulation correlates with developmental trends in mechanical recovery

To quantify viscoelastic properties we utilized a technique which has often been used to investigate biofilm material properties: rheometry (56–58). Due to the different qualitative responses to physical perturbation we observed between WT and Δ*P*_*s*_ biofilms, we first developed a rheological recovery assay to quantify biofilm response to deformation and determine if sporulation altered damage and recovery (**H3**) or had no effect (**H1, H2**). To compare the effect of the Δ*P*_*s*_ mutation to previously studied factors in biofilm rheology, we also performed experiments with the matrix knockouts Δ*pg*_*s*_*B* and Δ*ep*_*s*_ *A*-*O* (Materials and Methods). In this assay, we used a rheometer (Fig. 2A) to measure the storage modulus of a biofilm while applying high strain for short periods of time. This resulted in a time-series of biofilm viscoelastic response (Fig. 2BC), from which we extracted several metrics that could test possible hypotheses for how sporulation affects biofilm viscoelasticity (Table 2).

Our first metric is the initial damage, which is defined as the drop in storage modulus due to high strain normalized by the initial low-strain modulus (Fig. 2D). We used this metric to quantify the relative softening that the biofilms experienced when subjected to high strain. At 24 hours, none of the strains exhibited statistically significant differences in their initial damage. By 72 hours, the spore-knockout biofilms exhibited a statistically significant change in immediate damage, dropping relative to WT, while the WT and PGA-knockout biofilms showed no statistically significant differences in immediate damage. Furthermore, they exhibited no statistically significant differences from their 24 hour values, demonstrating recovery property consistency over time in spore-producing biofilms. The EPS-knockout showed a slight increase in immediate damage compared to WT at 72 hours. Despite the statistical significance, the small relative deviation from WT suggested that this difference was not biologically meaningful.

Our second recovery metric is the recovery fraction, defined as the rebound in storage modulus following high strain normalized by the drop in modulus due to high strain (Fig. 2E). We used this metric to quantify the transient changes in the polymer network: the greater the recovery fraction, the greater the fraction of the initial decrease in modulus was transient. At 24 hours, there was no statistically significant difference in recovery fraction between the strains. At 72 hours, only the spore-deficient biofilms showed a statistically significant difference in recovery, decreasing relative to WT. None of the spore-producing biofilms exhibited statistically significant differences in their means. However, all spore-producing biofilms increased in average recovery fraction from 24 to 72 hours.

The memory index, our final recovery metric, is defined as the final low-strain storage modulus (following the second period of high strain) normalized by the initial low-strain modulus (Fig. 2F). We used the memory index to quantify how well the biofilm recovers, or remembers, its initial storage modulus following multiple rounds of injuries, i.e., high-strain events. At 24 hours, none of the strains showed a statistically significant difference from WT. By 72 hours, the spore-deficient biofilms had an average memory index less than that of WT, while spore-producing biofilms had average indices close to WT. Spore-knockout biofilms had a mean memory index much less than their 24-hour value, while spore-producing biofilms had indices that increased over time.

In all three metrics, we observed a significant difference in viscoelastic response between Δ*P*_*s*_ and WT and matrix mutants, a finding inconsistent with the rigid inclusions (**H1**) and network density (**H2**) hypotheses but consistent with the hypothesis that sporulation alters biofilm polymer network restructuring processes (**H3**), leading to distinct damage and recovery (Table 2). Across our three metrics, the matrix and spore mutants showed no significant differences from WT at 24 hours. This suggests that the polymer network restructuring processes active at 24 hours were similar between biofilms. However, by 72 hours, the spore-deficient biofilms diverged from the matrix-mutant and WT biofilms, suggesting a divergence in polymer network restructuring mechanisms and yielding, consistent with **H3**. In an effort to characterize these mechanisms and further test our hypotheses, we investigated the stress relaxation spectra of the biofilms.

### Sporulation narrows and maintains biofilm relaxation spectrum

To assess the effect of sporulation on the biofilm stress relaxation spectrum, we collected data using a frequency sweep assay. Our hypotheses predicted no spectrum change under **H1**, spectrum broadening under **H2**, and spectrum narrowing under **H3**. To determine the spectrum, we fit the data to a generalized Maxwell model–the most general linear model of viscoelasticity–with three modes (Fig. 3A). We fixed the relaxation times to 0.01, 0.1, and 1.0 seconds and fit the mode moduli to the storage and loss moduli simultaneously (Fig. 3B). We normalized the mode moduli using the total modulus (*G*_*total*_ = *G*_*inf*_ + *G*_1_ + *G*_2_ + *G*_3_), resulting in the normalized moduli representing the fraction of the total modulus active at each time scale.

At 24 hours, only spore-knockout biofilms had a statistically significant deviation from WT in their mode moduli (Fig. 3C). The spore-knockout 0.01, 0.1, and 1.0 s average mode moduli were 55%, 54%, and 88% that of WT, respectively. The spore-knockout biofilms had an average *G*_*inf*_ 1.5 × WT.

At 72 hours, the only statistically significant deviation in average mode moduli was for the spore-knockout 0.01 second mode which was 80% that of WT (Fig. 3D). The spore-knockout average *G*_*inf*_ was 1.29× WT.

There was no significant difference in the mode moduli of the spore-producing biofilms between 24 and 72 hours (Fig. 3CD).

These data indicated that the long-term modulus *G*_*inf*_ was smaller in spore-producing biofilms, leading to a narrower relaxation spectrum consistent with **H3**. In order to more fully investigate this as well as biofilm viscoelastic properties in the linear regime, we next performed a strain sweep assay.

### Sporulating biofilms have equivalent linear properties

We measured the stiffness and viscosity of biofilms to further rule out possible mechanisms for the distinct mechanical properties of ΔP_*s*_ colonies (Table 2). Hypothesis **H1** predicted sporulation would increase stiffness, **H2** predicted it would decrease stiffness but increase viscosity, and **H3** predicted both stiffness and viscosity would decrease. To determine biofilm baseline viscoelastic properties, we measured the plateau storage and loss moduli (Fig. 4AB) and loss factor (Fig. 4C).

**Figure 4.**
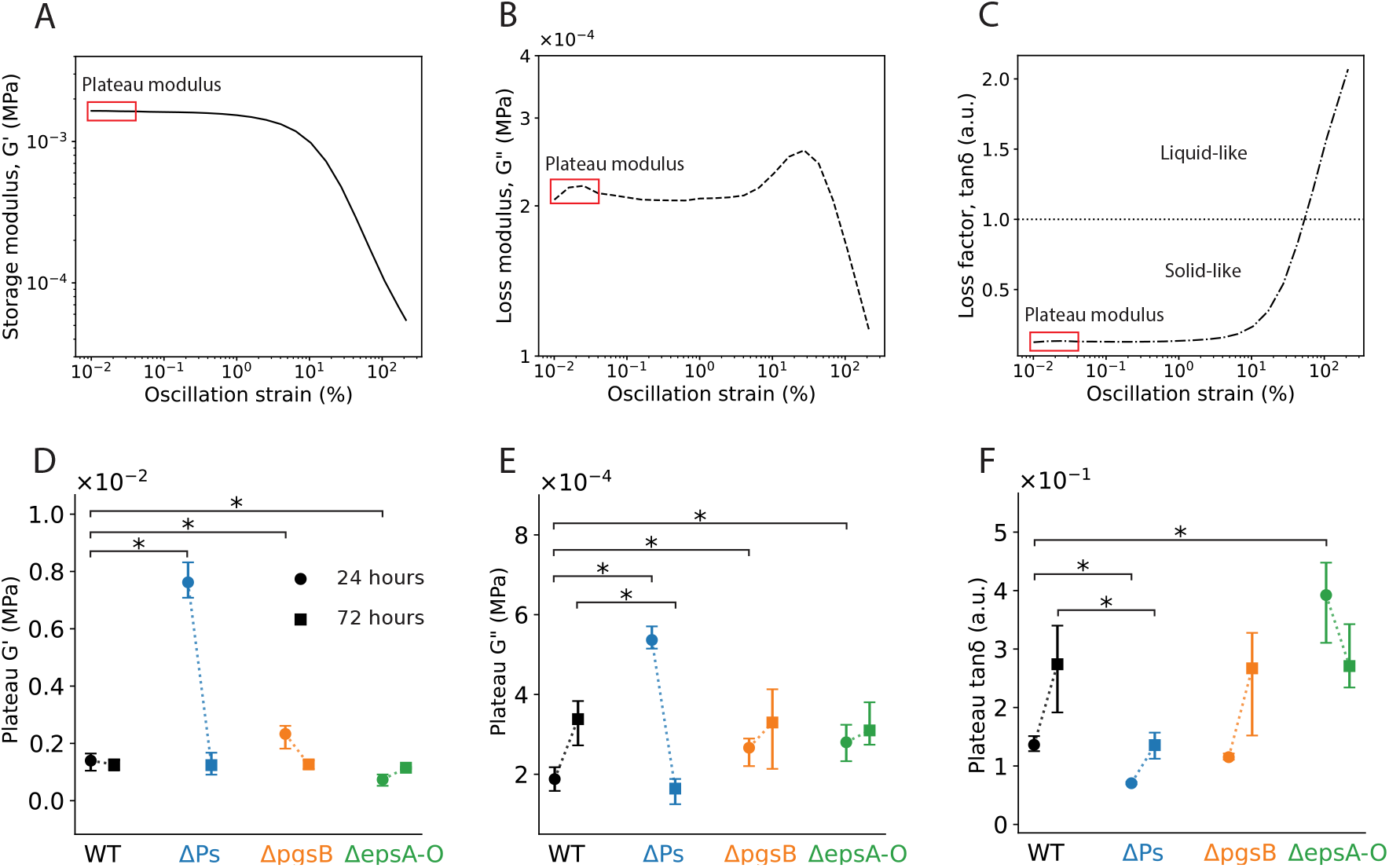
Sporulation maintains biofilm storage and loss moduli. (A) Storage modulus, (B) loss modulus, and (C) loss factor vs oscillation strain amplitude for a representative strain sweep dataset. The average of the boxed data constitutes the plateau value. (D) Average plateau storage modulus. The 24-hour spore-knockout biofilms have an average 4× that of WT. The PGA-knockout biofilms have an average slightly greater and the EPS-knockout biofilms slightly less than WT. There is no significant variation in the average storage modulus at 72 hours. (E) Average plateau loss modulus. At 24 hours, the mutant biofilms have average moduli greater than WT, with the spore-knockout biofilms exhibiting the greatest difference. At 72 hours, the WT and matrix-mutant biofilms converge to similar moduli while the spore-knockout biofilms drop significantly. (E) Average plateau loss factor. The spore-knockout biofilms have a smaller average loss factor compared to WT at 24 hours. At this time point, the EPS-knockout biofilms is greater in average loss factor than WT. At 72 hours, the spore-knockout biofilms are significantly smaller in average loss factor than WT. (D) - (F) Error bars represent 95% confidence intervals determined by the Margin of Error (ME = SEM × t^∗^). *p<0.05. P-values calculated by Welch’s two-tailed t-test on independent samples.

The storage moduli of the mutants all showed statistically significant differences from WT at 24 hours (Fig. 4D). The average storage modulus of the spore-knockout biofilms was 5 times that of WT. The PGA-knockout and EPS-knockout biofilms had average storage moduli 1.6 and 0.33 times that of WT, respectively. However, by 72 hours, none of the mutants had any significant difference from WT.

The loss moduli exhibited a similar trend, with all mutants showing a statistically significant difference from WT at 24 hours (Fig. 4E). The PGA-and EPS-knockout biofilms had loss moduli 1.4 times that of WT, while the spore-knockout biofilms had moduli 2.9 times that of WT. At 72 hours, only the spore-knockout biofilms showed a statistically significant difference, with an average 0.44 times that of WT.

At 24 hours, the spore-knockout biofilms had an average loss factor 0.43 times that of WT (Fig. 4F). The EPS-knockout biofilms had an average loss factor 2.86 times that of WT. The PGA-knockout biofilms had no significant difference from WT at this time point. At 72 hours, the only biofilms to exhibit a significant difference in loss factor were the spore-knockout biofilms, with an average 0.52 times that of WT.

These data are consistent with sporulation raising the biofilm noise temperature *x* (i.e., pushing the system towards a less glassy, more fluid-like state) and lowering the network density *ρ*, consistent with **H2** and **H3**. Furthermore, the data indicate that *x* and *ρ* change from 24 to 72 hours. We further explore this interpretation, as well as that of the data from preceding assays, in the following section.

## DISCUSSION

We have shown that sporulation affects biofilm physical properties in several important ways. Spore production is associated with greater damage and recovery at 72 hours, a finding in agreement with the network restructuring hypothesis, **H3**. Furthermore, these results contradict the predictions of the rigid inclusions (**H1**) and network density (**H2**) hypotheses. We have also shown that sporulation was correlated with a narrowing of the biofilm stress relaxation spectrum, in accordance with **H3** and contradiction to **H1** and **H2**. Finally, we saw that spore production was associated with a decrease in biofilm stiffness and viscosity at 24 hours, in agreement with **H3** and disagreement with **H1** and **H2**. However, by 72 hours the stiffness and viscosity shifted as can be understood by **H2** and **H3**. In this section, we discuss these findings in greater detail.

### Polymer network restructuring processes undergo divergent development in spore-producing biofilms

At 72 hours, the spore-knockout biofilms exhibit a drop in initial damage that is statistically and physically significant. This drop in immediate damage is consistent with a polymer network that has, compared to WT, a lower yield rate. This contrasts with the results at 24 hours, which showed no significant difference in the immediate damage between WT and spore-knockout biofilms, suggesting that the networks had similar yield rates.

These results suggest that spore-deficient biofilms have polymer networks with distinct yielding and restructuring processes from those of spore-producing biofilms, consistent with **H3**. That spore-deficient biofilms diverge from 24 to 72 hours, while spore-producing biofilms maintain their recovery metrics, suggests that spore-producing biofilms have network restructuring processes that are better-maintained over time, in contrast to spore-deficient biofilms. Although our recovery data are consistent with our network restructuring model, we cannot identify the specific molecular composition and processes of the biofilms, making molecular studies of intact biofilms an avenue ripe for future work.

### Stress response spectrum is maintained in spore-producing biofilms

In the frequency sweep assay, the fitted values of the mode moduli *G*_*i*_ are best interpreted as the strength of the biofilm’s stress relaxation at short (0.01 s), intermediate (0.1 s) and long (1.0 s) time scales. Although the generalized Maxwell model does not uniquely identify microscopic mechanisms, the 0.01 second mode is commonly interpreted as a local matrix relaxation process (59). The 0.1 second mode may be interpreted as the cooperative rearrangement of small connected regions of the matrix, rather than purely local relaxation or whole-network restructuring (60). The 1.0 second mode reflects persistent relaxation of longer-lived load-bearing structure, that is, stress redistribution requiring motion or reorganization over larger connected portions of the matrix (61). These results further demonstrate that determining the specific molecular mechanisms and how they correspond to relaxation at these time scales would be of additional value.

The normalized mode moduli *G*_*i*_/*G*_*tot*_ were smaller for all three modes in the spore-knockout biofilms at 24 hours compared to WT. This indicates that spore-knockout biofilms undergo less stress relaxation at the fitted time scales compared to WT, shifting a greater fraction of the stress to *G*_*inf*_ . That is to say, the spore-deficient biofilms retain a greater fraction of stress indefinitely, suggesting a polymer network with distinct restructuring processes, consistent with **H3**. This is further supported by the observed differences in yielding and recovery, as previously discussed, and greater magnitude storage modulus that the spore-knockout biofilms exhibit at 24 hours, as will be discussed below. By 72 hours, the spore-knockout’s relaxation spectrum advantage is less pronounced, with only the shortest relaxation modulus less than WT, and in turn a greater *G*_*inf*_ . This may be interpreted as the spore-knockout biofilms either decreasing in network density *ρ* or increasing in noise temperature *x* over time, in line with **H2** and **H3**, respectively. In contrast, the WT and matrix-mutant biofilms maintained consistent mode moduli from 24 to 72 hours, indicating greater stability in their stress relaxation processes over time. Taken together, these results indicate that by 72 hours the polymer network restructuring processes that are active in spore-knockout biofilms have become more similar to those active in spore-producing biofilms. That the relaxation spectrum changes significantly over time in spore-knockout biofilms but remains constant in spore-producing biofilms demonstrates that spore production alters both the relaxation processes in the biofilm and the way in which those processes change over time. This is evidence that spore production grants physical property stability over time.

### Stiffness is consistent but lesser in spore-producing biofilms

At 24 hours, all mutant biofilms exhibited a statistically significant deviation from WT in average storage modulus. However, only the spore-knockout biofilms exhibited an average storage modulus that was also significantly different in magnitude as well, being 5 times that of WT. This suggests that the spore-knockout biofilms are stiffer than WT and the other mutants. This increase in storage modulus is consistent with a more highly connected or mechanically intact network, possibly from a lack of cell lysis associated with sporulation or death at this time point, indicating a smaller noise temperature *x* in agreement with **H3**. This may be additionally understood as the spore-knockout biofilms having a greater network density *ρ* due to a greater fraction of the population producing matrix in line with **H2**.

At 72 hours, we observed that all biofilms had similar storage moduli. Specifically, the biofilms all converged to an average storage modulus value matching that of WT at 24 hours. However, the spore-producing biofilms increased in loss modulus, whereas the spore-knockout biofilms decreased. Taken together, these results are consistent with the spore-producing biofilms maintaining their noise temperature *x* while decreasing in network density *ρ* in line with **H2** and **H3**.

These results highlight that network properties and re-structuring processes differ heavily on the basis of spore production. This further supports that sporulation has a much more significant impact than PGA or EPS production alone on the establishment and development of biofilm rheological properties through its effect on network density *ρ* and noise temperature *x* as detailed in **H2** and **H3**.

## CONCLUSION

Our results demonstrate that sporulation, and the *spo0A* regulatory pathway that governs it, shapes not only the survival of individual *B. subtilis* cells but also the mechanical development trajectory of the biofilm community as a whole. Spore-deficient biofilms initially assemble a stiffer, more viscous polymer network than WT, reflected in elevated storage and loss moduli and a relaxation spectrum shifted toward longer timescales. This early mechanical advantage, however, is not sustained: by 72 hours, spore-deficient biofilms yield and recover more slowly, and their moduli and relaxation spectra converge toward WT values. Wild-type and matrix-mutant biofilms, by contrast, largely maintain stable viscoelastic properties, and even demonstrate greater recovery, across this same developmental window, indicating a polymer network that is developing more stably than and distinctly from spore-deficient biofilms.

These findings indicate that sporulation confers a distinct mechanical benefit at the population level: rather than simply protecting individual dormant cells, activation of the sporulation pathway may stabilize the biofilm matrix against the structural breakdown otherwise observed as the community matures. This links a well-characterized cell-fate decision to an emergent, community-scale material property, suggesting that gene regulatory programs traditionally understood in terms of single-cell survival can also function as determinants of collective mechanical resilience. Our findings demonstrate that spores acting as rigid inclusion (**H1**) or modifying the biofilm polymer network density (**H2**) are alone insufficient to explain the impact of sporulation on the establishment and development of viscoelastic properties. Rather, our findings indicate that sporulation alters both biofilm network density and restructuring processes, consistent with **H2** and **H3**.

This work provides a basis for identifying the specific matrix components and cellular processes (e.g., reduced lysis, altered EPS or PGA deposition, shifts in matrix entanglement and cross-linking, etc.) that are important in biofilm viscoelastic development in sporulating bacterial species. Therefore, it is clear that further work to understand the molecular basis of the bulk properties and dynamics we described here is needed and should be an avenue of future study. Identifying which matrix components sustain viscoelastic stability in spore-producing biofilms would allow us to distinguish network density from restructuring effects, to identify the mechanisms underpinning stress relaxation, and to target the properties that govern biofilm persistence and spore dispersal. The expanded understanding of this topic will not only aid in mechanically describing biofilms, but facilitate the development of techniques for their management, leading to improvements in industrial and medical outcomes.

## Supporting information

Spore- video

PGA- video

EPS- video

## DATA AVAILABILITY

All original data and code is available at https://github.com/Larkin-Lab/ducharme_larkin_sporulation.git and is publicly available as of the date of publication. Additional information is available upon request from the corresponding author.

## AUTHOR CONTRIBUTIONS

Conceptualization: N.A.D., J.W.L. Formal analysis: N.A.D. Funding acquisition: J.W.L. Investigation: N.A.D. Methodology: N.A.D. Project administration: J.W.L. Resources: N.A.D. Supervision: J.W.L. Visualization: N.A.D., J.W.L. Writing – original draft: N.A.D. Writing, review & editing: N.A.D., J.W.L.

## ACKNOWLEDGMENTS

This work was supported by NIH R35GM142584 (J.W.L.) and a Burroughs Wellcome Fund CASI award (J.W.L.). We extend special thanks to Dr. Abigail Plummer for discussions on theoretical models, and the members of the Larkin Lab for general feedback. We acknowledge the Boston University Biomedical Engineering Core Facilities for use of its TA Instruments DHR-2 Rheometer.

## SUPPORTING CITATIONS

References(55, 62–67) appear in the supporting material.

## SUPPORTING MATERIAL

An online supplement to this article can be accessed at https://github.com/Larkin-Lab/ducharme_larkin_sporulation.git.

## REFERENCES

1. Gloag, E. S., S. Fabbri, D. J. Wozniak, and P. Stoodley, 2020. Biofilm mechanics: Implications in infection and survival. Biofilm 2:100017.

2. Wells, M., R. Schneider, B. Bhattarai, H. Currie, B. Chavez, G. Christopher, K. Rumbaugh, and V. Gordon, 2023. Perspective: The viscoelastic properties of biofilm infections and mechanical interactions with phagocytic immune cells. Frontiers in Cellular and Infection Microbiology 13.

3. Charlton, S. G. V., M. A. White, S. Jana, L. E. Eland, P. G. Jayathilake, J. G. Burgess, J. Chen, A. Wipat, and T. P. Curtis, 2019. Regulating, Measuring, and Modeling the Viscoelasticity of Bacterial Biofilms. Journal of Bacteriology 201.

4. Peterson, B. W., Y. He, Y. Ren, A. Zerdoum, M. R. Libera, P. K. Sharma, A.-J. van Winkelhoff, D. Neut, P. Stoodley, H. C. van der Mei, and H. J. Busscher, 2015. Viscoelasticity of biofilms and their recalcitrance to mechanical and chemical challenges. FEMS Microbiology Reviews 39:234–245.

5. Kragh, K. N., J. B. Hutchison, G. Melaugh, C. Rodesney, A. E. L. Roberts, Y. Irie, P. Ø. Jensen, S. P. Diggle, R. J. Allen, V. Gordon, and T. Bjarnsholt, 2016. Role of Multicellular Aggregates in Biofilm Formation. mBio 7.

6. Rumbaugh, K. P., and K. Sauer, 2020. Biofilm dispersion. Nature Reviews Microbiology 18:571–586.

7. Gordon, V. D., and L. Wang, 2019. Bacterial mechanosensing: the force will be with you, always. Journal of Cell Science 132.

8. Rodesney, C. A., B. Roman, N. Dhamani, B. J. Cooley, P. Katira, A. Touhami, and V. D. Gordon, 2017. Mechanosensing of shear by Pseudomonas aeruginosa leads to increased levels of the cyclic-di-GMP signal initiating biofilm development. Proceedings of the National Academy of Sciences 114:5906–5911.

9. Zhang, Q., D. Nguyen, J. B. Tai, X. Xu, J. Nijjer, X. Huang, Y. Li, and J. Yan, 2022. Mechanical Resilience of Biofilms toward Environmental Perturbations Mediated by Extra-cellular Matrix. Advanced Functional Materials 32.

10. Hindieh, P., J. Yaghi, J. C. Assaf, A. Chokr, A. Atoui, N. Tzenios, N. Louka, and A. E. Khoury, 2025. Emerging Multimodal Strategies for Bacterial Biofilm Eradication: A Comprehensive Review. Microorganisms 13:2796.

11. Adeboye, A., H. Onyeaka, Z. Al-Sharify, and N. Nnaji, 2024. Understanding the Influence of Rheology on Biofilm Adhesion and Implication for Food Safety. International Journal of Food Science 2024.

12. Tallawi, M., M. Opitz, and O. Lieleg, 2017. Modulation of themechanical propertiesof bacterial biofilmsinresponse to environmental challenges. Biomaterials Science5:887–900.

13. Secchi, E., 2026. Biofilms as living soft materials: Towards a mechanistic description of their linear and nonlinear mechanics. Phys. Rev. E 114:021001.

14. Pandit, S., M. Fazilati, K. Gaska, A. Derouiche, T. Nypelö, I. Mijakovic, and R. Kádár, 2020. The Exo-Polysaccharide Component of Extracellular Matrix is Essential for the Viscoelastic Properties of Bacillus subtilis Biofilms. International Journal of Molecular Sciences 21:6755.

15. Rana, S., and L. S. B. Upadhyay, 2020. Microbial exopolysaccharides: Synthesis pathways, types and their commercial applications. International Journal of Biological Macromolecules 157:577–583.

16. Burla, F., Y. Mulla, B. E. Vos, A. Aufderhorst-Roberts, and G. H. Koenderink, 2019. From mechanical resilience to active material properties in biopolymer networks. Nature Reviews Physics 1:249–263.

17. Chew, S. C., B. Kundukad, T. Seviour, J. R. C. van der Maarel, L. Yang, S. A. Rice, P. Doyle, and S. Kjelleberg, 2014. Dynamic Remodeling of Microbial Biofilms by Functionally Distinct Exopolysaccharides. mBio 5.

18. Charlton, S. G., A. N. Bible, E. Secchi, J. L. Morrell-Falvey, S. T. Retterer, T. P. Curtis, J. Chen, and S. Jana, 2023. Microstructural and Rheological Transitions in Bacterial Biofilms. Advanced Science 10.

19. Asally, M., M. Kittisopikul, P. Rué, Y. Du, Z. Hu, T. Çağatay, A. B. Robinson, H. Lu, J. Garcia-Ojalvo, and G. M. Süel, 2012. Localized cell death focuses mechanical forces during 3D patterning in abiofilm. Proceedings of the National Academy of Sciences 109:18891–18896.

20. Grobas, I., M. Polin, and M. Asally, 2021. Swarming bacteria undergo localized dynamic phase transition to form stress-induced biofilms. eLife 10.

21. Yuan, L., B. Zhang, H. Dai, Y. Miao, Z. Xu, Z. Yang, and X. an Jiao, 2025. Starvation-induced metabolic adaptation of Bacillus cereus during biofilm formation: Phenotypic characterization and proteomic analysis. Food Research International 220:117167.

22. Grossman, A. D., and R. Losick, 1988. Extracellular control of spore formation in Bacillus subtilis. Proceedings of the National Academy of Sciences 85:4369–4373.

23. Grossman, A. D., 1995. GENETIC NETWORKS CONTROLLING THE INITIATION OF SPORULATION AND THE DEVELOPMENT OF GENETIC COMPETENCE IN BACILLUS SUBTILIS. Annual Review of Genetics 29:477–508.

24. Hutchison, E. A., D. A. Miller, and E. R. Angert, 2014. Sporulation in Bacteria: Beyond the Standard Model. Microbiology Spectrum 2.

25. Majed, R., C. Faille, M. Kallassy, and M. Gohar, 2016. Bacillus cereus biofilms—same, only different. Frontiers in microbiology 7:1054.

26. Dürre, P., 2011. Ancestral sporulation initiation. Molecular microbiology 80:584–587.

27. Fujita, M., J. E. González-Pastor, and R. Losick, 2005. High-and low-threshold genes in the Spo0A regulon of Bacillus subtilis. Journal of bacteriology 187:1357–1368.

28. Huang, Q., Z. Zhang, Q. Liu, F. Liu, Y. Liu, J. Zhang, and G. Wang, 2021. SpoVG is an important regulator of sporulation and affects biofilm formation by regulating Spo0A transcription in Bacillus cereus 0–9. BMC microbiology 21:172.

29. Dapa, T., and M. Unnikrishnan, 2013. Biofilm formation by Clostridium difficile. Gut microbes 4:397–402.

30. Nishikawa, M., and K. Kobayashi, 2021. Calcium prevents biofilm dispersion in Bacillus subtilis. Journal of bacteriology 203:10–1128.

31. Ulrich, N., K. Nagler, M. Laue, C. S. Cockell, P. Setlow, and R. Moeller, 2018. Experimental studies addressing the longevity of Bacillus subtilis spores – The first data from a 500-year experiment. PLOSONE 13:e0208425.

32. Tan, I. S., and K. S. Ramamurthi, 2014. Spore formation in Bacillus subtilis. Environmental Microbiology Reports 6:212–225.

33. Turnbull, L., M. Toyofuku, A. L. Hynen, M. Kurosawa, G. Pessi, N. K. Petty, S. R. Osvath, G. Cárcamo-Oyarce, E. S. Gloag, R. Shimoni, U. Omasits, S. Ito, X. Yap, L. G. Monahan, R. Cavaliere, C. H. Ahrens, I. G. Charles, N. Nomura, L. Eberl, and C. B. Whitchurch, 2016. Explosive cell lysis as a mechanism for the biogenesis of bacterial membrane vesicles and biofilms. Nature Communications 7:11220.

34. Vlamakis, H., Y. Chai, P. Beauregard, R. Losick, and R. Kolter, 2013. Sticking together: building a biofilm the Bacillus subtilis way. Nature Reviews Microbiology 11:157–168.

35. Arnaouteli, S., N. C. Bamford, N. R. Stanley-Wall, and Ákos T. Kovács, 2021. Bacillus subtilis biofilm formation and social interactions. Nature Reviews Microbiology 19:600–614.

36. Mielich-Süss, B., and D. Lopez, 2015. Molecular mechanisms involved in Bacillus subtilis biofilm formation. Environmental Microbiology 17:555–565.

37. Dogsa, I., B. Bellich, M. Blaznik, C. Lagatolla, N. Ravenscroft, R. Rizzo, D. Stopar, and P. Cescutti, 2024. Bacillus subtilis EpsA-O: A novel exopolysaccharide structure acting as an efficient adhesive in biofilms. npj Biofilms and Microbiomes 10:98.

38. Rajput, R., S. Saha, H. S. Gill, M. Rajeev, and S. Pandit, 2025. Microbially derived (W-PGA) poly-W-glutamic acid: applications and commercial uses across industries. Preparative Biochemistry & Biotechnology 1–27.

39. Branda, S. S., F. Chu, D. B. Kearns, R. Losick, and R. Kolter, 2006. A major protein component of the Bacillus subtilis biofilm matrix. Molecular microbiology 59:1229–1238.

40. Saha, A., J. M. Jones, A. Plummer, and J. W. Larkin, 2026. Formation of a swelling gel underlies a morphological transition in Bacillus subtilis biofilms. bioRxiv .

41. Konkol, K. Blair, 2013. Plasmid-Encoded ComI Inhibits Competence in the Ancestral 3610 Strain of Bacillus subtilis. Journal of Bacteriology 195:4085–4093.

42. Jones, J. M., M. Yao, A. Mugler, and J. W. Larkin, 2026. Colony morphogenesis regulates sporulation dynamics in bacterial biofilms. bioRxiv .

43. Jones, J. M., I. Grinberg, A. Eldar, and A. D. Grossman, 2021. A mobile genetic element increases bacterial host fitness by manipulating development. Elife 10:e65924.

44. Branda, S. S., J. E. González-Pastor, S. Ben-Yehuda, R. Losick, and R. Kolter, 2001. Fruiting body formation by Bacillussubtilis. Proceedingsof theNational Academy of Sciences 98:11621–11626.

45. Alger, M. S. M., 1997. Polymer science dictionary, Chapman & Hall, 218 & 287.

46. Abdalla, H. M. A., 2025. Review of rules of mixture for effective elastic properties in fibrous and particulate composite materials. Composite Structures 367:119216.

47. Winter, H. H., and F. Chambon, 1986. Analysis of Linear Viscoelasticity of a Crosslinking Polymer at the Gel Point. Journal of Rheology 30:367–382.

48. Mours, M., and H. H. Winter, 1996. Relaxation Patterns of Nearly Critical Gels. Macromolecules 29:7221–7229.

49. Stauffer, D., A. Coniglio, and M. Adam, 1982. Gelation and critical phenomena, Springer Berlin Heidelberg, 103–158.

50. Sollich, P., 1998. Rheological constitutive equation for a model of soft glassy materials. Physical Review E 58:738–759.

51. Wingender, J., T. R. Neu, and H.-C. Flemmings, 1999. Microbial Extracellular Polymeric Substances. Springer Berlin Heidelberg.

52. Fei, C., S. Mao, J. Yan, R. Alert, H. A. Stone, B. L. Bassler, N. S. Wingreen, and A. Košmrlj, 2020. Nonuniform growth and surface friction determine bacterial biofilm morphology on soft substrates. Proceedings of the National Academy of Sciences 117:7622–7632.

53. Vlamakis, H., C. Aguilar, R. Losick, and R. Kolter, 2008. Control of cell fate by the formation of an architecturally complex bacterial community. Genes & development 22:945–953.

54. van Oosten, A. S. G., X. Chen, L. Chin, K. Cruz, A. E. Patteson, K. Pogoda, V. B. Shenoy, and P. A. Janmey, 2019. Emergence of tissue-like mechanics from fibrous networks confined by close-packed cells. Nature 573:96–101.

55. Sollich, P., F. Lequeux, P. Hébraud, and M. E. Cates, 1997. Rheology of Soft Glassy Materials. Physical Review Letters 78:2020–2023.

56. Gordon, V. D., M. Davis-Fields, K. Kovach, and C. A. Rodesney, 2017. Biofilms and mechanics: a review of experimental techniques and findings. Journal of Physics D: Applied Physics 50:223002.

57. Lieleg, O., M. Caldara, R. Baumgärtel, and K. Ribbeck, 2011. Mechanical robustnessof Pseudomonasaeruginosa biofilms. Soft matter 7:3307–3314.

58. Asp, M. E., M.-T. Ho Thanh, D. A. Germann, R. J. Carroll, A. Franceski, R. D. Welch, A. Gopinath, and A. E. Patteson, 2022. Spreading rates of bacterial colonies depend on substrate stiffness and permeability. PNAS Nexus 1.

59. Johari, G. P., and M. Goldstein, 1970. Viscous Liquids and the Glass Transition. II. Secondary Relaxations in Glasses of Rigid Molecules. The Journal of Chemical Physics 53:2372–2388.

60. Adam, G., and J. H. Gibbs, 1965. On the Temperature Dependence of Cooperative Relaxation Properties in Glass-Forming Liquids. TheJournal of Chemical Physics 43:139–146.

61. Rubinstein, M., and A. N. Semenov, 2001. Dynamics of Entangled Solutions of Associating Polymers. Macromolecules 34:1058–1068.

62. Liu, Z.-L., and X. Chen, 2022. Water-Content-Dependent Morphologies and Mechanical Properties of Bacillus subtilis Spores’ Cortex Peptidoglycan. ACSBiomaterials Science & Engineering 8:5094–5100.

63. Kesel, S., S. Grumbein, I. Gümperlein, M. Tallawi, A.-K. Marel, O. Lieleg, and M. Opitz, 2016. Direct Comparison of Physical Properties of Bacillus subtilis NCIB 3610 and B-1 Biofilms. Applied and Environmental Microbiology 82:2424–2432.

64. Rubinstein, M., and R. H. Colby, 2018. Polymer Physics, Oxford University Press, 253–281.

65. Rogers, S. S., C. van der Walle, and T. A. Waigh, 2008. Microrheology of Bacterial Biofilms In Vitro: Staphylo coccus aureus and Pseudomonas aeruginosa. Langmuir 24:13549–13555.

66. Pavlovsky, L., J. G. Younger, and M. J. Solomon, 2013. In situ rheology of Staphylococcus epidermidis bacterial biofilms. Soft Matter 9:122–131.

67. Pasqui, D., M. De Cagna, and R. Barbucci, 2012. Polysaccharide-Based Hydrogels: The Key Role ofWater in Affecting Mechanical Properties. Polymers 4:1517–1534

