## Supplemental Information for "Sporulation modulates viscoelastic development through extracellular matrix restructuring in *Bacillus subtilis* biofilms"

#### 1. VISCOELASTIC HYPOTHESES

Here, we apply simple models of material properties to predict the physical impact of sporulation on biofilms. Each model addresses a hypothesis for the effect of sporulation. The first hypothesis, H1, is that spores act as rigid inclusions (i.e., particles interspersed throughout a polymer network that bolster its stiffness). We address H1 with the rule of mixtures (see Section 1A). The second hypothesis, H2, is that sporulation reduces the number of vegetative cells which in turn reduces the quantity and density of extracellular matrix. We use gelation theory to address H2 (see Section 1B). The third hypothesis, H3, is that sporulation causes the polymer network to restructure more easily (i.e., for the processes influencing cross-links, entanglements, and other network parameters to have a more active role). We address H3 using soft glassy rheology.

##### A. Rule of mixtures

In order to make predictions based on H1 (that spores act as rigid inclusions) we utilize the rule of mixtures. The rule of mixtures is used in materials science to estimate upper- and lower-bounds for the physical properties, including stiffness, of composite materials. Here, we treat vegetative cells and extracellular matrix as one material and spores as another, making the biofilm a composite material consisting of cells + matrix and spores. We group vegetative cells and matrix in a distinct group from spores due to the similarity in their independent elastic moduli and because their moduli are many orders of magnitude smaller than that of spores (1, 2).

$$G^* = fG_1^* + (1-f)G_2^* \tag{S1}$$

$$G^* = \left(\frac{f}{G_1^*} + \frac{1-f}{G_2^*}\right)^{-1} \tag{S2}$$

For a composite material composed of two constituents, the composite complex modulus $G^*$ is predicted by Eqs. S1 and S2, where $G_1^*$ and $G_2^*$ are the complex moduli of the constituents and $f$ is the volume fraction of the first constituent. Equation S1, the rule of mixtures, is derived from the Voigt model and assumes that the strain is equal in both constituents. This equation sets the upper bound on the composite modulus. Equation S2, the inverse rule of mixtures, is derived from the Reuss model and assumes that the stress is equal in both constituents. This equation sets the lower bound on the composite modulus. Here, we note that real materials must fall somewhere between the Voigt and Reuss models (i.e., somewhere between having constituent materials with equal strain or stress) although few materials of interest are likely purely Voigt- or Reuss-like. This allows for upper and lower bounds to be placed on composite material properties.

Both Eqs. S1 and S2 predict an increase in $G^*$ with an increase in $f$. Additionally, both predict increases in $G'$ and $G''$, the storage (elastic) and loss (viscous) moduli, with an increase in $f$. In the case of bacterial biofilms, $f$ would represent the volume fraction of spores and $1 - f$ the volume fraction of everything else (i.e., the vegetative cells and extracellular matrix). This would mean that as the spore fraction increases, so too would the composite modulus, with greater spore $G'$ and $G''$ leading to a more pronounced effect. Spores are known to have a greater elastic modulus than bulk matrix (approximately $10^6\times$) (1, 2), so spore fraction would have a large effect on the composite $G'$. In contrast, spores do not have a well-defined independent viscosity, making their contribution to the composite $G''$ minimal. For these reasons, H1 predicts that knocking out sporulation ($\Delta P_s$) would decrease stiffness. However, our data show the opposite trend, with moduli decreasing in spore-producing biofilms, indicating that H1 is insufficient to explain the role of sporulation in biofilm material properties.

##### A.1. Derivations

Here we derive Eqs. S1 and S2.

$$\gamma = \gamma_1 = \gamma_2 \tag{S3}$$
$$\sigma = G^* \gamma \tag{S4}$$

In the Voigt model, we assume the constituents have equal strain (Eq. S3) and apply Hooke's law (Eq. S4).

$$\sigma = f\sigma_1 + (1 - f)\sigma_2 \tag{S5}$$
$$G^*\gamma = fG_1^*\gamma_1 + (1 - f)G_2^*\gamma_2 \tag{S6}$$
$$G^* = fG_1^* + (1 - f)G_2^* \tag{S7}$$

With forces balanced at the interface between materials, we arrive at Eq. S5, which when combined with Eqs. S3 and S4 yields Eq. S7, which is identical to Eq. S1.

$$\sigma = \sigma_1 = \sigma_2 \tag{S8}$$

In the Reuss model, we assume the constituents have equal stress (Eq. S8) and utilize Hooke's law (Eq. S4).

$$\gamma = f\gamma_1 + (1-f)\gamma_2 \tag{S9}$$

$$\frac{\sigma}{G^*} = \frac{f\sigma_1}{G_1^*} + \frac{(1-f)\sigma_2}{G_2^*} \tag{S10}$$

$$\frac{1}{G^*} = \frac{f}{G_1^*} + \frac{1-f}{G_2^*} \tag{S11}$$

$$G^* = \left(\frac{f}{G_1^*} + \frac{1-f}{G_2^*}\right)^{-1} \tag{S12}$$

Strain distributes between the constituents according to Eq. S9, which when combined with Eqs. S8 and S4 yields Eq. S12, which is identical to Eq. S2.

See *Polymer Science Dictionary* by Alger (3) and Abdalla (4) for additional information.

##### B. Networks in gelation regime

Hypothesis 2 predicts that sporulation's primary effect on biofilm rheological properties is to reduce the density of the colony's matrix network: if more cells become dormant spores, fewer cells are in a state a matrix production, resulting in a less dense polymer network. To predict what effect this would have on rheological measurements, we must relate network density to measured moduli, a pursuit to which percolation theory lends itself well. This approach is defensible since *Bacillus subtilis* biofilms are known to be near the sol-gel percolation point (5).

$$G \approx \frac{\rho RT}{M_x} + T_e G_e \tag{S13}$$

Percolation theory predicts that, for a polymer network in the gelation regime, elastic modulus scales with network density, as described by Eq. S13 (6), where $G$ is the elastic modulus, $\rho$ the network density, $R$ the gas constant, $T$ the temperature, $M_x$ the number-average apparent molar mass of a network strand, $T_e$ the entanglement trapping factor, and $G_e$ the entanglement modulus.

Equation S13 predicts that elastic modulus $G$ decreases with a decrease in network density $\rho$. In the case of biofilms, having more spores may correlate with having a lower density of extracellular matrix (compared to comparable biofilms without spores). Therefore, according to Eq. S13, spore-producing biofilms would have a lower elastic modulus.

$$\epsilon = \frac{\rho - \rho_c}{\rho_c} \tag{S14}$$

$$G \sim \epsilon^t \tag{S15}$$

Assuming that biofilm polymer networks are not far from the percolation point (as demonstrated in (5)), it is convenient use the parameter $\epsilon$ as defined in Eq. S14 (7) to write an expression for $G$ as in Eq. S15 (7). The parameter $\rho$ here represents the network density and $\rho_c$ the gelation critical density. The scaling exponent $t$ (not time) dictates how $G$ changes with respect to $\epsilon$.

$$\eta \sim \epsilon^{-s} \tag{S16}$$

Percolation theory allows us to write a similar expression for the dynamic viscosity $\eta$: Eq. S16 (7). Here $s$ is a scaling exponent representing how strongly network density lowers viscosity.

From Eqs. S14 - S16 we see that a decrease in network density $\rho$ corresponds to a decrease in $G$ and an increase in $\eta$.

$$H(\tau) \sim \tau^{-n} \text{ for } \tau \ll \tau_z \tag{S17}$$

In the gelation regime, percolation theory also predicts that the relaxation spectrum follows Eq. S17 (8). Here, $H(\tau)$ represents the relaxation spectrum, $\tau$ is the relaxation time, $n$ is a scaling exponent, and $\tau_z$ is the relaxation spectrum cutoff time. Above the gelation point, the theory predicts a stable long-term elastic contribution as $\tau$ approaches infinity: $G_{inf}$.

$$\tau_z \sim |\epsilon|^{-\Delta} \tag{S18}$$

The cutoff relaxation time $\tau_z$ is defined by Eq. S18 (9) where $\epsilon$ is the distance from the critical gel point density $\rho_c$ and $\Delta$ is a class scaling exponent. The quantity $\epsilon$ varies with network density according to Eq. S14.

From Eqs. S17 - S14 we see that a decrease in network density $\rho$, as from sporulation, results in an increase in the cutoff relaxation time $\tau_z$ leading to a broader relaxation spectrum $H(\tau)$. Additionally, a looser relaxation spectrum may be interpreted as a more disordered network.

##### C. Soft glassy rheology

Soft glassy rheology (SGR) is a phenomenological model of material properties (10, 11). This model consists of mesoscopic elements that transition between energy "traps" in response to strain.

$$E' = E - \frac{1}{2}kl^2 \tag{S19}$$

$$\rho(E') \sim e^{-E'} \tag{S20}$$

$$P_{eq}(E') \sim \rho(E')e^{-E'/x} \tag{S21}$$

The depth of the traps (defined by Eq. S19) determines the strain $l$ necessary for an element to escape the trap and fall into one of different energy. The energy landscape of these traps is denoted by $\rho(E')$ and takes the form of exponential decay as shown in Eq. S20. In the soft glassy fluid regime, which is the phase most applicable to biofilms (12–14), the material's probability distribution evolves towards the Boltzmann distribution as defined by Eq. S21.

The parameter $x$, known as the "noise temperature", is a mean-field subsumption of the processes through which elements transition between traps. The transition of elements between traps is what we refer to as network restructuring, although in reality it likely includes this and other processes. The noise temperature is key in determining if the material behaves like a true glass ($x < 1$), a soft glassy fluid ($1 < x < 2$), an ordinary Newtonian/Maxwell-like fluid ($x = 2$), or a simple liquid ($x > 2$).

$$G'(\omega) \sim G'_0 \left(\frac{\omega}{\omega_0}\right)^{x-1} \tag{S22}$$

$$G''(\omega) \sim G''_0 \left(\frac{\omega}{\omega_0}\right)^{x-1} \tag{S23}$$

Under SGR (in the soft glassy regime: $1 < x < 2$), the shear moduli take the form of Eqs. S22 and S23.

$$P_{eq}(\tau) \sim \tau^{-x} \tag{S24}$$

Additionally, the relaxation time spectrum exhibits power law behavior as described by Eq. S24 for $\tau \geq 0.1$ s.

We propose that sporulation allows easier element rearrangement (i.e., network restructuring, etc.), increasing $x$. With an increase in $x$, Eqs. S22 and S23 predict a decrease in $G'$ and $G''$, a trend qualitatively in agreement with our data. Additionally, an increase in $x$ results in faster 'decay' in Eq. S24, resulting in a narrower relaxation spectrum, consistent with our observations.

$$\Gamma \sim \Gamma_0 e^{-E'/x} \tag{S25}$$

Finally, the damage and recovery metrics laid out in the main text can also be explained through SGR. SGR describes damage as elements having sufficient strain to push them into different energy traps, resulting in yielding at a rate defined by Eq. S25.

As $x$ increases, as we propose in the case of sporulation, Eq. S25 predicts an increase in the element yield rate, corresponding to an increase in damage (as defined in the main text). Predictions for recovery are less definitive under SGR: SGR predicts that the noise temperature $x$ affects the recovery rate, but it doesn't make a definitive prediction of whether it increases or decreases recovery. Applying SGR in detail to further describe biofilm properties and predict the impact of sporulation is an avenue of future work.

#### 2. SUPPLEMENTAL DATA

All data and analysis code can be accessed at http://www.biophysj.org and https://github.com/Larkin-Lab/ducharme_larkin_sporulation.git.

##### A. CFU counts and spore percentages

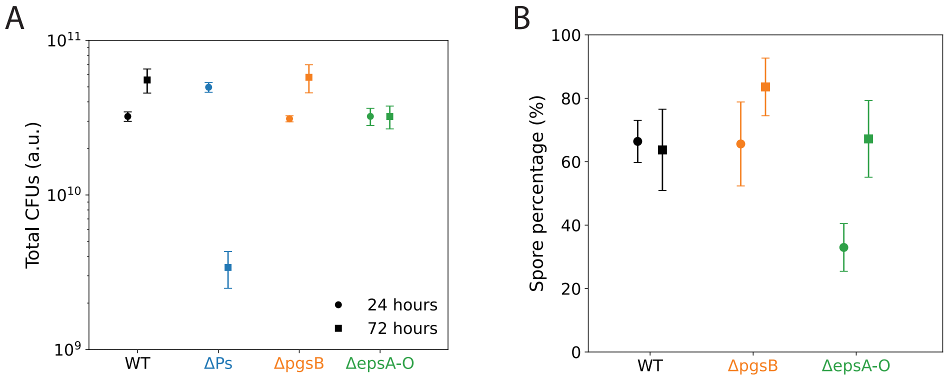


**Fig. S1.** (A) Total CFUs: arithmetic mean and standard deviation. At 24 hours, the spore-knockout has greater total CFUs than WT (1.66×). At 72 hours, the spore-knockout has CFUs much less than 72-hour WT (0.064×) and 24-hour spore-knockout (0.07×). Wild-type and the PGA-knockout have similar total CFUs at both 24 and 72 hours, with both showing an increase with time. The EPS-knockout is constant in total CFUs from 24 to 72 hours. (B) Spore percentages: geometric mean and standard deviation. The EPS-knockout has a lower spore percentage at 24 hours compared to WT (0.77×). At 72 hours, the PGA-knockout has a higher spore percentage than WT (0.37×). N = 3 independent samples per strain.

$$N_{\text{CFUs}} = N_{\text{counts}} * 10^{n+2} \tag{S26}$$

We calculated the number of CFUs using Eq. S26, where $n$ is the power of the dilution (6 or 7) and the factor of $10^2$ accounts for the initial and plated volumes of 10 mL and 100 $\mu$L, respectively.

Figure S1 shows CFU counts and spore percentages for the bacterial strains used in this work. The spore-knockout has greater total CFUs than WT at 24 hours. The PGA- and EPS-knockouts have comparable total CFUs to WT at 24 hours. At 72 hours, the spore-knockout drops in total CFUs substantially. The EPS-knockout has slightly lower total CFUs than WT at that time point.

The WT and PGA-knockout biofilms have similar spore percentages at 24 hours, and the EPS-knockout has a lower percentage. At 72 hours, the PGA-knockout has a higher spore percentage than WT whereas the EPS-knockout has a comparable percentage to WT.

These data indicate it is unlikely that the total CFUs or spore percentage explain any rheological differences of the PGA- and EPS-knockouts from WT. However, it is possible that these quantities have some indirect impact on rheological properties. In particular the EPS-knockout, which exhibits some differences in these quantities, may partially contribute to the minor differences in rheological properties noted in the main text. The spore-knockout also has substantial differences in both total CFUs and spore percentage compared to WT which may indirectly contribute to differences in rheological properties.

#### B. Mass measurements

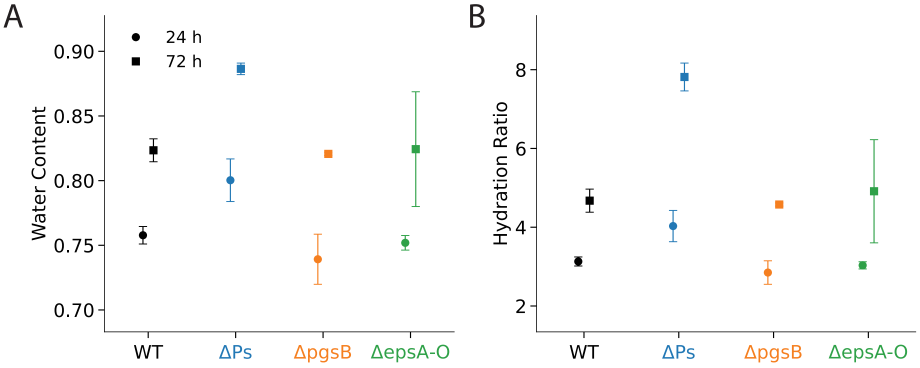


**Fig. S2.** Plots of biofilm (A) water content and (B) hydration ratio at 24 and 72 hours. The water content is the ratio of bound water mass to wet mass. The hydration ratio is the ratio of bound water mass to dry mass. At 24 and 72 hours, spore-knockout biofilms have greater average water content and hydration ratio compared to spore-producing biofilms (WT, matrix-knockouts). All biofilms increase in both metrics from 24 to 72 hours. N = 3 independent samples per strain.

$$m_{bound} = m_{wet} - m_{dry} \tag{S27}$$
$$WC = \frac{m_{bound}}{m_{wet}} \tag{S28}$$
$$HR = \frac{m_{bound}}{m_{dry}} \tag{S29}$$

We calculated the mass of bound water in the biofilm as the difference between the wet and dry masses of the biofilm (Eq. S27). We took the mass of the extracellular matrix, cells, and spores as the dry mass. We defined the water content (WC) as the ratio of the bound water mass to the wet, or total, mass (Eq. S28) and the hydration ratio (HR) as the ratio of bound water mass to dry mass (Eq. S29).

Figure S2 shows the bound water content and hydration ratio of biofilms formed by the bacterial strains used in this work. Both metrics are greater in the spore-knockout biofilms ($\Delta P_s$) than the spore-producing biofilms (WT, $\Delta pgsB$, $\Delta epsA$-$O$). Also, both metrics increase from 24 to 72 hours.

These data suggest bound water variations from sporulation may play some role in biofilm rheological properties. The data show that there is more bound water in spore-knockout biofilms. However, having more bound water generally decreases both the storage and loss moduli of gels and gel-like materials (15). Because we find that the storage modulus is actually greater in the spore-knockout at 24 hours (and shows no variation from WT at 72 hours) (see Fig. S7), bound water is insufficient to explain our rheological data.

#### C. Recovery assay data

Figure S3 shows the mean storage modulus vs time with standard deviations for our four bacterial strains at 24 and 72 hours. At 24 hours, the PGA- and EPS-knockout biofilms exhibited an upward drift during the low-strain periods, possibly indicating that these biofilms have not yet structural equilibrium. This data is further discussed in the main text.

#### D. Frequency sweep fits

Figure S4 shows representative fits and relative residuals using generalized Maxwells models with one-, two-, and three-modes. The one-mode model fits a mode at 0.01 s and $G_{inf}$ and the two mode-model additionally fits a mode at 0.1 s. The one- and two-mode fits are of poor quality. The three-mode Maxwell model, which fits an additional mode at 1.0 s, is the lowest-order model for which the fits are acceptable.

Figures S5 and S6 show the mean, standard deviations, fits, and relative residuals of the three-mode generalized Maxwell model with our frequency sweep data. The fits and relative residuals are generally of

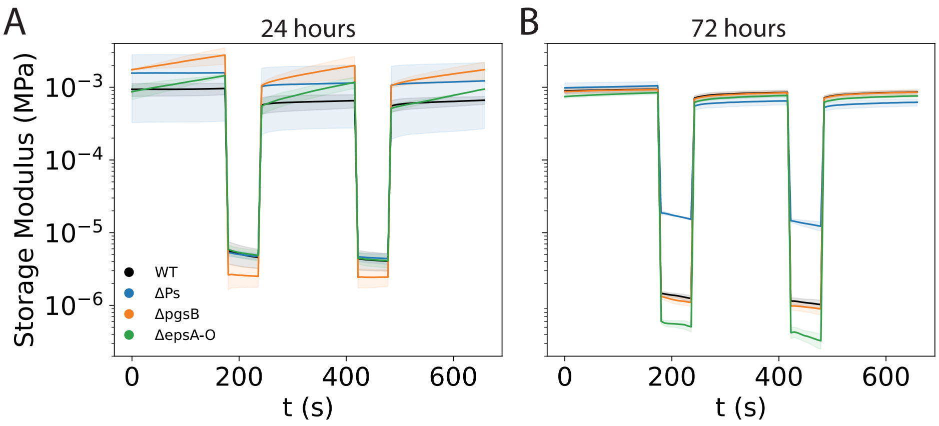


**Fig. S3.** Mean storage modulus vs time at (A) 24 and (B) 72 hours measured with alternating periods of low- and high-amplitude strain. Shaded regions indicate standard deviations. Both panels share the same y-limits. N = 3 independent samples per strain.

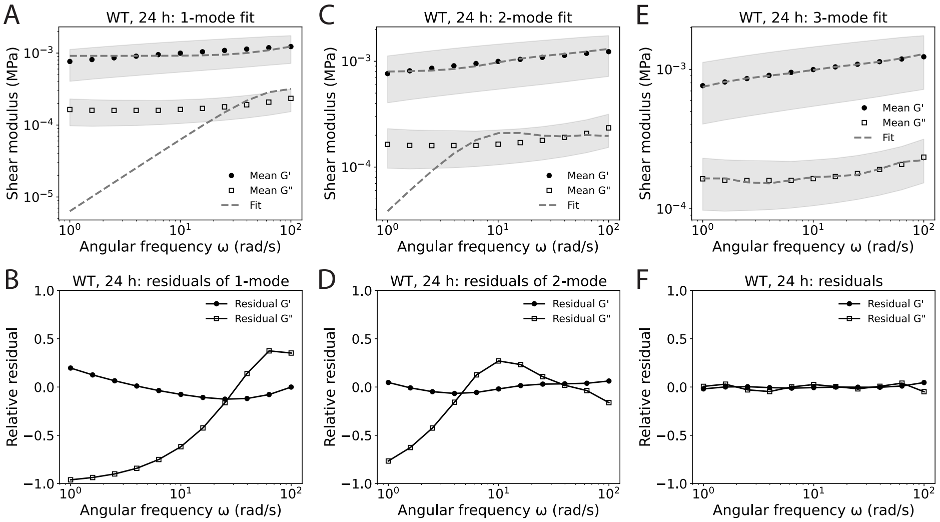


**Fig. S4.** Multi-mode Maxwell fits and relative residuals. Average fit and relative residuals for one- (A, B), two- (C, D), and three-mode (E, F) generalized Maxwell models to WT storage and loss modulus data at 24 hours. The relative residuals for the one- (B) and two-mode (D) models are large, and much larger than that of the three-mode (F) model. Additionally, the lower-order fits (A, C) are qualitatively much poorer than that of the three-mode (E) model. N = 3 independent samples.

good quality. Each replicate was fit separately. The fitting parameters were averaged and from the averages the fits were plotted.

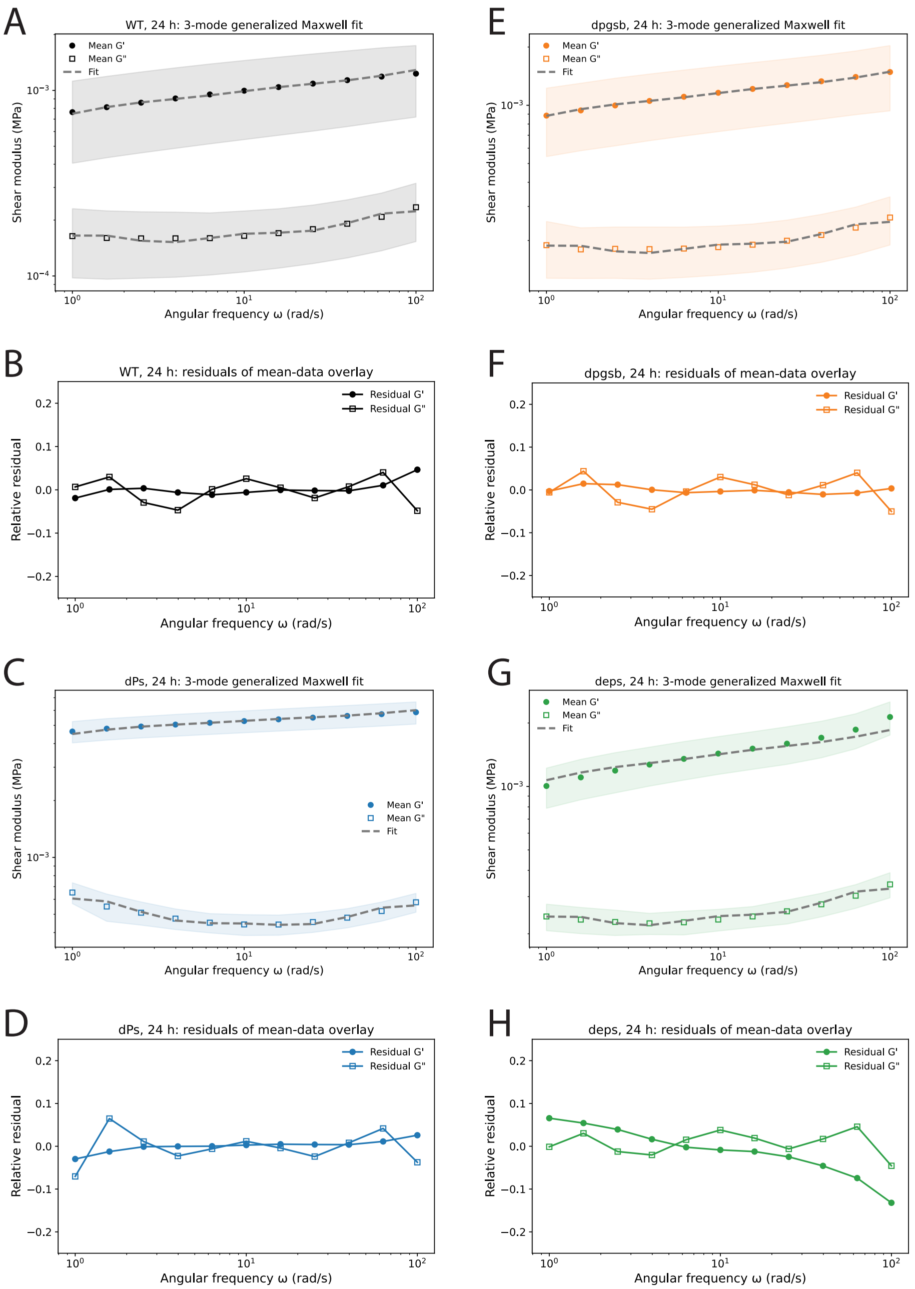


**Fig. S5.** Mean storage and loss moduli with fit to three-mode generalized Maxwell model at 24 hours for (A) WT, (C) $\Delta P_s$, (E) $\Delta pgsB$, and (G) $\Delta epsA$-$O$ biofilms. Relative fit residuals (residual divided by modulus) for (A) WT, (C) $\Delta P_s$, (E) $\Delta pgsB$, and (G) $\Delta epsA$-$O$ biofilms. Shaded regions indicate standard deviations. N = 3 independent samples per strain.

#### E. Strain sweep data

Figure S7 shows the mean storage and loss moduli and loss factor vs strain amplitude with the shaded regions showing the standard devi-

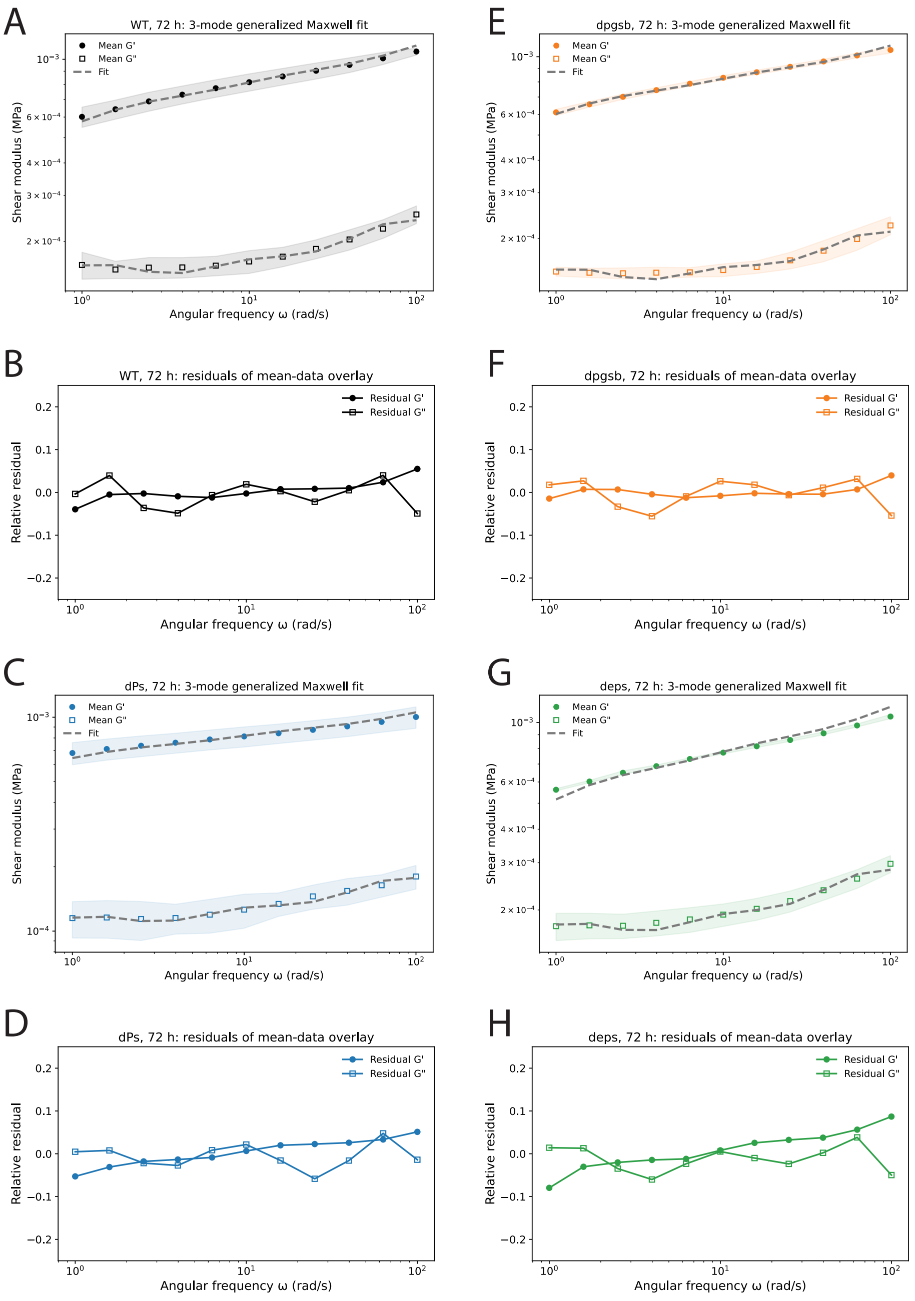


**Fig. S6.** Mean storage and loss moduli versus angular frequency with fit to three-mode generalized Maxwell model at 72 hours for (A) WT, (C) $\Delta P_s$, (E) $\Delta$*pgsB*, and (G) $\Delta$*epsA-O*biofilms. Relative fit residuals (residual divided by modulus) for (A) WT, (C) $\Delta P_s$, (E) $\Delta$*pgsB*, and (G) $\Delta$*epsA-O* biofilms. Shaded regions indicate standard deviations. N = 3 independent samples per strain.

ations. Plateau moduli were extracted from these data, the analysis of which is described in the main text.

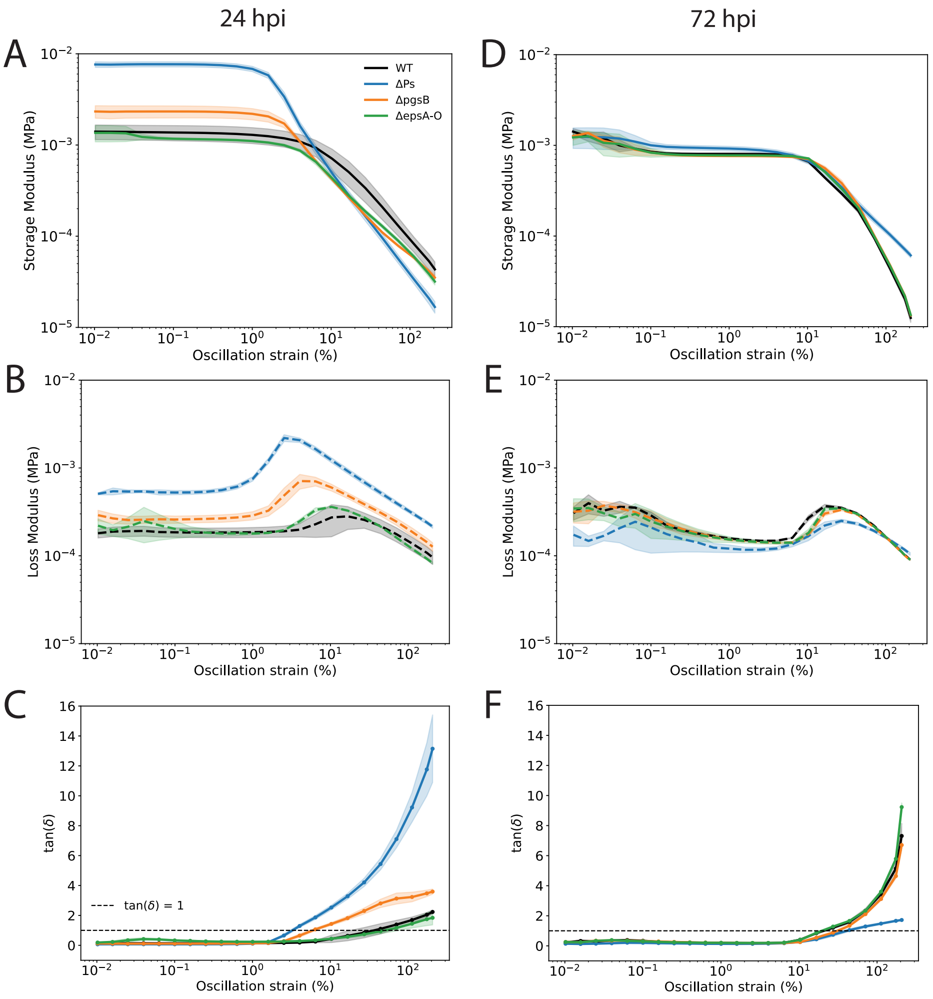


**Fig. S7.** Mean storage moduli vs strain amplitude at (A) 24 and (D) 72 hours. Mean loss moduli at (B) 24 and (E) 72 hours. Mean loss factor at (C) 24 and (F) 72 hours. Shaded regions indicate standard deviations. N = 3 independent samples per strain.

#### F. Stereo microscope videos of biofilm scraping

We recorded videos using a stereo microscope as described in the main text. These videos are accessible with the other supplemental materials at http://www.biophysj.org and at https://github.com/Larkin-Lab/ducharme_larkin_sporulation.git.

The videos were recorded with the same exposure settings so any differences in apparent brightness are due to physical differences between biofilms.

We dragged the bottom of a micro-centrifuge tube through the center of the biofilms, displacing a small volume of biomass. We then recorded videos over the following three minutes. We observed a difference in the recoil of the biofilms based on their ability to sporulate. The wild-type (spore-producing) biofilm exhibited greater recoil, i.e., greater return towards its pre-injured shape, compared to the spore-knockout biofilms. This observation hinted at a difference in the rheological properties of the biofilms based on sporulation ability, which led to the research presented here.

### REFERENCES


1. Liu, Z.-L., and X. Chen, 2022. Water-Content-Dependent Morphologies and Mechanical Properties of *Bacillus subtilis* Spores' Cortex Peptidoglycan. *ACS Biomaterials Science & Engineering* 8:5094–5100.

2. Kesel, S., S. Grumbein, I. Gümperlein, M. Tallawi, A.-K. Marel, O. Lieleg, and M. Opitz, 2016. Direct Comparison of Physical Properties of Bacillus subtilis NCIB 3610 and B-1 Biofilms. *Applied and Environmental Microbiology* 82:2424–2432.

3. Alger, M. S. M., 1997. Polymer science dictionary, Chapman & Hall, 218 & 287.

4. Abdalla, H. M. A., 2025. Review of rules of mixture for effective elastic properties in fibrous and particulate composite materials. *Composite Structures* 367:119216.

5. Saha, A., J. M. Jones, A. Plummer, and J. W. Larkin, 2026. Formation of a swelling gel underlies a morphological transition in *Bacillus subtilis* biofilms. *bioRxiv* .

6. Rubinstein, M., and R. H. Colby, 2018. Polymer Physics, Oxford University Press, 253–281.

7. Stauffer, D., A. Coniglio, and M. Adam, 1982. Gelation and critical phenomena, Springer Berlin Heidelberg, 103–158.

8. Winter, H. H., and F. Chambon, 1986. Analysis of Linear Viscoelasticity of a Crosslinking Polymer at the Gel Point. *Journal of Rheology* 30:367–382.

9. Mours, M., and H. H. Winter, 1996. Relaxation Patterns of Nearly Critical Gels. *Macromolecules* 29:7221–7229.

10. Sollich, P., F. Lequeux, P. Hébraud, and M. E. Cates, 1997. Rheology of Soft Glassy Materials. *Physical Review Letters* 78:2020–2023.

11. Sollich, P., 1998. Rheological constitutive equation for a model of soft glassy materials. *Physical Review E* 58:738–759.

12. Rogers, S. S., C. van der Walle, and T. A. Waigh, 2008. Microrheology of Bacterial Biofilms In Vitro: *Staphylococcus aureus* and *Pseudomonas aeruginosa*. *Langmuir* 24:13549–13555.

13. Pavlovsky, L., J. G. Younger, and M. J. Solomon, 2013. In situ rheology of Staphylococcus epidermidis bacterial biofilms. *Soft Matter* 9:122–131.

14. Charlton, S. G. V., M. A. White, S. Jana, L. E. Eland, P. G. Jayathilake, J. G. Burgess, J. Chen, A. Wipat, and T. P. Curtis, 2019. Regulating, Measuring, and Modeling the Viscoelasticity of Bacterial Biofilms. *Journal of Bacteriology* 201.

15. Pasqui, D., M. De Cagna, and R. Barbucci, 2012. Polysaccharide-Based Hydrogels: The Key Role of Water in Affecting Mechanical Properties. *Polymers* 4:1517–1534.
